# *In vivo* imaging uncovers an abundant but rarely active pool of plant ARP2/3 and its link to exocytosis and endocytosis

**DOI:** 10.64898/2026.07.10.737699

**Authors:** Barbora Jelínková, Maria Voloshina, Klára Liebezeit, Jana Krtková, Judith García-González, Stanislav Vosolsobě, Adam Harmanec, Kateřina Malínská, Ayoub Stelate, Eva Kollárová, Anežka Baquero Forero, Viktor Žárský, Kateřina Schwarzerová

## Abstract

- The ARP2/3 complex nucleates branched actin networks, but its spatiotemporal organization and cellular functions in plants remain poorly understood. We investigated the dynamics and cellular context of the ARP2/3–WAVE/SCAR machinery in living Arabidopsis cells.
- We combined high-resolution microscopy techniques with genetic, colocalization and co-immunoprecipitation analyses. Automated particle tracking quantified the dynamics and abundance of cortical ARP2/3 foci and their association with cytoskeletal and membrane-trafficking markers.
- ARP2/3 and WAVE/SCAR subunits formed abundant, highly dynamic cortical foci with an approximate dwell time of 1 second. Genetic and colocalization analyses indicated that most foci represent partially assembled or inactive complexes. Microtubules influenced foci abundance, whereas actin primarily affected their dynamics. ARP2/3 associated with both endocytic and exocytic markers. Exocytosis monitored by EXO84b was functionally altered in ARP2/3 mutants, while the dynamics of ARP2/3 remained unaffected in exo84b mutant. ARP2/3 subunits also co-immunoprecipitated with the exocyst subunit EXO84b.
- Our findings support a model in which plant cells maintain a large cortical reserve of ARP2/3 complexes that is only locally active. The association with both actin filaments and microtubules, and with exocytosis and endocytosis suggest that ARP2/3 contributes to cytoplasmic membrane remodeling dynamics and potentially links the cytoskeleton, membrane trafficking and cell surface organization.

## Introduction

Plants employ two known mechanisms of regulated actin nucleation: the extensive and functionally very redundant formin family, and the actin related protein 2/3 (ARP2/3) complex (Cvrčková et al., 2004; D. Szymanski & Staiger, 2017). The ARP2/3 is an eukaryotic conserved heteroheptameric complex, which nucleates branched actin filaments that give rise to a fine actin meshwork (Volkmann et al., 2001). The name-giving subunits ARP2 and ARP3 are structurally related to monomeric actin (Robinson et al., 2001); upon transient activation by a nucleation promoting factor (NPF), ARP2/3 complex binds to an existing actin filament, while ARP2 and ARP3 subunits change conformation to act as a seed for nucleation of a new daughter actin filament (Volkmann et al., 2001). The remaining five subunits, ARPC1-5, are unique proteins that provide interactions of the complex with the actin filaments during nucleation (Rouiller et al., 2008). While eukaryotic cells use a wide range of NPFs to activate ARP2/3, plants encode and utilize only the WAVE/SCAR complex (Yanagisawa et al., 2013). The activating complex WAVE/SCAR consists of 5 subunits, some of which are encoded by multiple paralogs in Arabidopsis - SCAR1-4, NAP1, BRICK1, ABIL1-4 and PIR1 (D. B. Szymanski, 2005). Mutants with non-functional NPF or ARP2/3 complexes have similar and non-additive phenotypes, implying that both complexes function in the same pathways (Basu et al., 2005; Yanagisawa et al., 2013). Although viable and fertile, plants with knocked-out genes encoding subunits of either ARP2/3 or WAVE/SCAR exhibit phenotypes that would seriously compromise survival under natural environmental conditions. Known defects point to the cytoskeleton—membrane—cell wall continuum being affected: distorted trichomes, changed epidermal cell morphology, fragmented vacuole and epidermal adhesion defects (reviewed in Jelínková et al., 2026).

Despite the fact that the mechanism of ARP2/3 activation and nucleation of a new actin filament from a daughter filament, leading to the formation of specific branched actin, is very well understood at the molecular level, the function of this specific actin structure in plants remains unclear. Recent advanced image analysis suggests that the ARP2/3 complex is not involved in the global reorganization of actin in most analyzed plant cells (Cifrová et al., 2020; García-González et al., 2026; Xu et al., 2024), and dynamic branched actin is presumed to have highly localized roles (Yanagisawa et al., 2015). The mechanism of function of ARP2/3 complex-nucleated actin, which leads to the phenotypes described above in mutant plants, is very difficult to study due to the high dynamics of plant actin and its high concentration in cells. Therefore, high-resolution spatiotemporal analysis of the localization and dynamics of the ARP2/3 complex and its NPFs may provide mechanistic insight into the cellular functions of ARP2/3-mediated actin nucleation in plant cells.

The assembled ARP2/3 complex is a relatively small structure, measuring only 10-15 nm in size (Robinson et al., 2001), and it is expected to be quite dynamic, mirroring the rapid rearrangement of branched actin arrays (Pollard & Borisy, 2003). Thus, under these conditions, visualization in living cells can be expected to be difficult and fixed-cell imaging might not sufficiently reflect its spatial organization and dynamics within the cell. Nevertheless, earlier studies visualizing ARP2/3 in plants using immunolocalization put the complex in proximity to actin filaments, microtubules, and organelles. The antibodies often detected structures that decorated the filaments (Fišerová et al., 2006; Van Gestel et al., 2003; C. Zhang, Mallery, & Szymanski, 2013) and also appeared as a dispersed granular pattern, distinct punctate structures of varying sizes, or a large nucleus-associated rod-like object (C. Zhang, Mallery, & Szymanski, 2013). These observations are consistent across a variety of cell types - tobacco leaves and maize roots (Van Gestel et al., 2003), Arabidopsis cotyledons and leaves (C. Zhang, Mallery, & Szymanski, 2013), or BY-2 cell culture (Fišerová et al., 2006; Havelková et al., 2015). Similarly, the subunits of WAVE/SCAR displayed a granular and reticulate pattern in the cytoplasm, more pronounced puncta of varying sizes colocalizing with endoplasmic reticulum (ER) and also plasma membrane (PM) adjacent accumulation in the three-way cell junctions (C. Zhang, Mallery, & Szymanski, 2013). Collectively, the immunolocalization experiments suggested that the ARP2/3 and WAVE/SCAR complexes are relatively abundant in the cytoplasm of plant cells and form several functionally distinct populations.

*In vivo* imaging confirmed the punctate nature of ARP2/3 and WAVE/SCAR localization, however, many of the described structures appear substantially larger than expected for individual 10-15 nm ARP2/3 complexes, indicating that these signals represent localized accumulations rather than single complexes. Epifluorescence microscopy of RFP-ARP3 in BY-2 cells, for example, revealed discrete puncta associated with actin nucleation sites (Maisch et al., 2009), whose size is apparently too large to represent single ARP2/3 complexes. Indeed, a number of works point to ARP2/3 or WAVE/SCAR accumulating in specific cellular domains. Fluorescent protein fusions of the WAVE/SCAR or ARP2/3 complexes subunits were observed to localize and accumulate to larger PM-adjacent domains - to three-way cell junctions, to developing lobes of cotyledon pavement cells (Dyachok et al., 2008; Qin et al., 2021), to peroxisomes (Martinek et al., 2023) and to sites subjected to mechanical pressure in epidermal cells associated with autophagy regulation at ER-PM contact sites (Wang et al., 2016, 2019) and during penetrative infection by the fungus *Blumeria graminis* (Qin et al., 2021). These localization patterns place the complexes at sites of active cell shaping, cell-cell contact or response to stress. WAVE/SCAR and ARP2/3 complex subunits can also be observed at the growing tip of Arabidopsis trichomes. This represents an important site of localization, as trichomes developing in the absence of a functional complex fail to achieve their wild type–like morphology, indicating an active role of ARP2/3 during growth (Mathur, 2005). Importantly, the structures observed in the growing tip of Arabidopsis trichomes, based on their dynamics, size, and close association with actin, demonstrate that they represent an *in vivo* observation of an individual dynamic actin-nucleating complex (Yanagisawa et al., 2015, 2018). In our previous work we demonstrated that a population of ARP2/3 and WAVE/SCAR accumulates in plant peroxisomes to facilitate pexophagy (Martinek et al., 2023). The size of these peroxisomal domains indicates the presence of multiple complexes and thus represents another example of a site within the plant cell where the WAVE/SCAR–ARP2/3 module accumulates. However, the high rate at which peroxisomes move prevented detailed analysis of the dynamics of this population. This work, together with our earlier observations (Havelková et al., 2015), revealed that, in addition to distinct accumulations, a substantial cytoplasmic signal is also detectable in cells expressing WAVE/SCAR and ARP2/3 markers.

In the present study, we aimed to characterize this cytoplasmic signal. We have employed variable angle epifluorescence microscopy (VAEM), which allowed us to observe protein complexes *in vivo* within the cortex, an optically accessible and spatially well-resolved cellular region. By combining multiple subunit visualization in mutant plants with image analysis, we demonstrate that the majority of ARP2/3 and WAVE/SCAR assemblies are short-lived, incomplete, and often inactive complexes. The demonstrated association with both actin and microtubules, as well as with endo- and exocytotic events, highlights its role at the nexus of cytoskeletal and membrane trafficking processes. Our results particularly emphasized the link between ARP2/3 and exocytosis. Beyond biological insights, our methodological approach provides a foundation for studying fast, noisy, and dynamic processes in plant cells.

## Materials and Methods

### Plant material and cultivation

*A. thaliana* genotypes used in this study were Col-0 (wt), *arpc5* (SALK_123936.4) (Cifrová et al., 2020), *arpc2a/distorted2-1* (El-Din El-Assal et al., 2004), *nap1* (SALK_038799) (Brembu et al., 2004; Li et al., 2004), *brk1-1* (Le et al., 2006), and *exo84b-1* (Fendrych et al., 2010). For *in vitro* cultivation, seeds were briefly washed with 96% ethanol and then surface sterilized by bleach solution (sodium hypochlorite solution at a final concentration of 2.5%) with 0.01% Triton X-100 for 10 min, washed in sterile H_2_O and sown on vertical plates with agar medium (1% (w/v) sucrose, 2.2 g l^−1^ Murashige and Skoog salts (Sigma-Aldrich, M5524) and 0.8% plant agar (Duchefa, P1001), pH 5.7) in a cultivation room under a photoperiod of 16 h light:8 h darkness, 23 °C and light intensity of 110 µmol m^−2^ s^−1^. For the expression of constructs with the β-estradiol-inducible system XVE, the medium was supplemented with β-estradiol (Sigma-Aldrich, E2758, stock solution 10 mM in DMSO). For cytoskeletal drug treatments, *in vitro*-grown 5-day-old (5 days after sowing, DAS) seedlings were transferred to liquid cultivation media supplemented with either 2 μM latrunculin B (LatB; Sigma-Aldrich, 428020, stock solution 2.57 mM in DMSO) to destabilize the actin filaments, 20 μM oryzalin (Ory; Sigma-Aldrich, PS410, stock solution 20 mM in DMSO) to destabilize the MTs, or an equivalent volume of DMSO as the mock treatment and incubated for 2 h before imaging. For the CK666 (Sigma-Aldrich, SML0006, stock solution 10 mM in DMSO) and pentabromopseudilin (PBP; MedChemExpress, Cat. No.: HY-113604, stock solution 10 mM in DMSO) treatments, plants were submerged in liquid cultivation media supplemented with 10 µM inhibitor or with equivalent volume of DMSO as a mock (Xu et al., 2024). For 2,3-butanedione monoxime (BDM, stock solution 500 mM in water) treatment, plants were submerged in liquid media supplemented with 50 mM BDM for 30 minutes with respective mock control.

### *In vivo* markers and molecular cloning

The *35S::mSc-ARPC5* construct was created using the GoldenBraid 2.0 system (Sarrion-Perdigones et al., 2013). First, the ARPC5 CDS was amplified from vector DNA (*35S::GFP-AtARPC5*, Martinek et al., 2023). For primers see Table S1. The PCR product was then ligated into a domestication pUPD2 vector using BsmBI and T4 Ligase. The construct was assembled into a transcription unit together with the 35S promoter, mScarlet-I and the *35S* terminator using the alpha 1 backbone vector. The transcription unit was then moved into the binary pDGB3omega1 vector with BsmBI (ThermoFisher) and BsaI (NEB) restriction enzymes and T4 Ligase (NEB). The *UBQ::GFP-AtARP5* was generated using the Gateway® cloning system (Invitrogen™) according to the protocol provided by the manufacturer., and was introduced into the empty vector, pDONOR P2 P3R, using the BP Clonase II (Invitrogen™). The resulting entry vector was recombined along with constitutive UBQ 10 promoter in pDONOR P4P1R (Kollárová et al., 2025), pEN L1-F-L2 (Karimi et al., 2007) and pH7m34GW (Karimi et al., 2005) using LR Clonase II (Invitrogen™) to create the final expression vector. All entry vectors were verified by sequencing using the M13. The *XVE::GFP* was produced by restriction cloning-genomic sequence for smRSGFP was amplified and cloned into an estrogen-inducible XVE vector (J. Zuo et al., 2000). The following fluorescent lines were also used: 35S::GFP-*At*ARPC5, 35S::*At*ARPC5-mCh, 35S::GFP-*At*ARPC2 (Martinek et al., 2023); XVE::GFP-*Nt*ARPC2 (Havelková et al., 2015); pNAP1::NAP1-GFP (Wang et al., 2016); pEXO84b::EXO84b-GFP (Fendrych et al., 2010); 35S::EXO84b-RFP (Fendrych et al., 2013); pLAT52::TPLATE-GFP (Van Damme et al., 2006), 35S::TPLATE-mRFP (Van Damme et al., 2011); pDRP1C:DRP1C-mOra (Konopka & Bednarek, 2008a), 35S::LifeAct-RFP (Fendrych et al., 2013); 35S::mCh-TUA5 (Gutierrez et al., 2009); ER-RFP marker (Nelson et al., 2007); 35S::GFP.

### Transformation of plant material

Arabidopsis plants were stably transformed by *Agrobacterium tumefaciens* strain C58C1 or GV3101 using the floral dip method (X. Zhang et al., 2006). Transformed seedlings were selected on agar plates with antibiotics.

### Microscopy and spectroscopy

#### VAEM

VAEM was used for imaging of the cortical cytoplasm of hypocotyl or cotyledon epidermal cells. Using 5-day-old seedlings of plants expressing the respective markers, roots and cotyledons were cut from the hypocotyls to ensure good contact between the tissue and the cover glass, and plants were observed immediately. A TIRF Zeiss Elyra SP.1 microscope equipped with an sCMOS PCO Edge 5.5 camera and with an alpha Plan-Apochromat 100×/1.46 oil objective was used for imaging cortical regions of hypocotyl epidermal cells with Highly Inclined and Laminated Optical sheet (HILO) mode. Zeiss Immersol 518 F for 30 °C was used as immersion media (RI 1.518). GFP (excitation 488 nm) was observed using a filter set with emission filter 495–550/LP 750. For colocalization analysis, GFP and RFP were excited by 488 and 561 nm, respectively, and a filter set with emission filter LBF 488/561 was used. Localization and colocalization were performed in time-lapse images (50-200 frames at 0.4 s and 0.2 s frame intervals, respectively). ZEN Blue software was used for image acquisition.

For dual localization of the cytoskeleton or ER with ARP2/3 foci, the microscope Elyra SP.7 with DuoLink camera was used. The alpha Plan-Apochromat 100×/1.46 oil or the alpha Plan-Apochromat 63×/1.46 oil objectives were used for imaging cortical regions of hypocotyl epidermal cells with HILO mode. The GFP (excitation 488 nm) and RFP or mCherry (excitation 561 nm) were observed in parallel using two pco.edge sCMOS (version 4.2 CL HS) cameras with the emission filter LBF 405/488/561/642 and dual camera beam splitter SBS LP 560. Time-lapse images were recorded at 50-200 frames at 0.2 s frame intervals. ZEN Blue software was used for image acquisition.

#### LSCM

To visualize GFP-NAP1 marker on the periclinal cell walls, whole 5-day-old seedlings were mounted in water and epidermal cells of cotyledons were imaged. The MIRAVA Polyscope (Abberior) equipped with the UPLSAPO60XW (1.2 NA, water immersion) objective and avalanche photodiode (APD) detectors was used. The GFP (excitation 488 nm) was detected within 498 nm - 647 nm wavelength window. RAYSHAPE adaptive optics was used for correction of optical distortions and data were deconvolved using Lightbox software. 3D reconstructions were created using the Arrivis 4D software.

#### Spinning disk

A spinning disk microscope Nikon (Eclipse Ti-E, inverted) equipped with a Photometrics Prime BSI camera and Plan-Apochromat L 100x/1.45 Oil objective was used to image EXO84b-GFP and TPLATE-GFP in the cortical cytoplasm of hypocotyls of 5-day-old seedlings. The GFP (excitation 488 nm) was observed using a filter set Semrock brightline Em 525/30. The exposure time was set to 400 ms and the interval between frames was 500 ms.

#### FRAP

FRAP was performed using a laser scanning confocal microscope Leica SP8 equipped with HC PL APO CS2 63×/1.2 W objective, using excitation 488 nm and emission 490– 550 nm (GFP). The pinhole was set to 1 AU at 580 nm. In cotyledon of 4-day-old seedling of the line *nap1*/pNAP1::NAP1-GFP, the site containing two to three three-way cell junctions was first imaged (pre-bleach). Subsequently, one 3-cell-junction together with part of the surrounding cytoplasm was bleached using the FRAP acquisition mode (120 s, laser power 100%) and imaged again (post-bleach). The site was further imaged after 5, 15, and 30 minutes. For the recovery analysis, mean signal intensities at sites of signal accumulation in the 3-cell-junctions ROI and in the surrounding cytoplasm ROI were compared between the bleached and non-bleached (reference) regions, using the vacuolar area within a single image as a reference background. The rate of recovery (F (t) *norm*) in 3-cell-junctions or in the cytoplasm ROIs was calculated using the following equation:

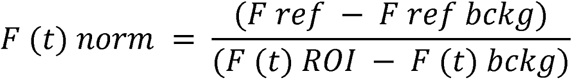

where *F ref* is the mean intensity measured before bleaching at the reference ROI, the *F ref bckg* is the background intensity measured before bleaching, and the *F(t) ROI* and *F (t) bckg* are intensities measured in the specific timepoint in measured ROI and in the background after bleaching, respectively. Further, the signal intensity ratio between the three-way junction and the adjacent cytoplasm in bleached and reference ROIs was calculated for each time point (including the pre-bleach) by dividing the mean fluorescence intensity values for either the bleached or non-bleached (reference) three-way junction (with the respective background values subtracted) by the adjacent cytoplasm mean fluorescence intensity values (with the respective background values subtracted). These values allowed to assess the degree of signal accumulation at the 3-cell-junction relative to the surrounding cytoplasm.

#### FCS

Arabidopsis seedlings were grown for 4 days on ½ MS medium supplemented with different β-estradiol concentrations for induction of gene expression. The analyzed lines were dis2-1/XVE::ARPC2-GFP and Col-0/XVE::GFP. Fluorescence correlation spectroscopy (FCS) data was acquired at anticlinal walls of cotyledon epidermal cells on a ZEISS LSM 900 laser scanning microscope (Axio Observer 7, inverted) equipped with Airyscan 2 Multiplex mode, using a C-Apochromat 63/1.20 W Korr UV VIS IR FCS objective. Data acquisition was performed using the FCS module Dynamics profiler (ZEISS). To ensure comparability across analyzed samples, the same conditions were used for all datasets.

### Protein isolation and co-IP

For total protein fraction isolation, approximately 1 g of 10-day-old Arabidopsis seedlings were frozen in liquid nitrogen and homogenized with a mortar and pestle. The frozen powder was mixed 1:1 (v/w) with 2× concentrated MES buffer (25 mM MES, 5 mM ethyleneglycol bis(2-aminoethyl ether)-N,N,N′,N′ tetraacetic acid, 5 mM MgCl_2_ and 1 M glycerol, pH 6.9) supplemented with a protease inhibitors cocktail (Sigma-Aldrich, P9599) and was let thaw on ice. The sample was then centrifuged at 4000 g for 20 min, followed by centrifugation of supernatant at 30000 g for 30 minutes, both at 4 °C. The supernatant was used for co-IP (input). 1 ml of this protein extract was mixed with 50 µl of magnetic beads from the µMACSTM GFP Isolation Kit (Miltenyi Biotec), and the mixture was incubated on ice for 1 h with mild shaking. The extract was then loaded into the column in a magnetic stand and GFP-tagged proteins and their interactors were isolated according to the manufacturer’s protocol. Isolated immunoprecipitated proteins (the whole amount of eluted fraction, 50 µl) as well as respective inputs (20 µl of extracts corresponding to 50 µg of total proteins) were separated using sodium dodecyl sulfate–polyacrylamide gel electrophoresis and transferred onto nitrocellulose membrane by electro-blotting (SemiDry, BioRad). Western blots were probed with rabbit anti-GFP (1:6000; Agrisera AS152987), rabbit anti-mCherry to detect RFP (1:6000; Abcam, ab167453), and respective secondary HRP-conjugated antibodies (goat anti-rabbit HRP conjugate ENZO ADI-SAB-300-J 1:10000) Proteins were visualized using the enhanced chemiluminescence (ECL) method (Pierce Western Blot-ting Substrate) using manual processing of X-ray films or Azure 600 Imaging System.

### Image analysis

#### Dwell time analysis

Time series images of length 50-200 frames were analyzed in Fiji (Schindelin et al., 2012). If needed, Bleach correction plugin was used to correct signal bleaching (Miura, 2020). A ROI was selected excluding any structures that would hinder the detection and tracking of foci (e.g. autofluorescence of chloroplast, peroxisome-associated ARP2/3 accumulations). Using the TrackMate plugin (Ershov et al., 2022; Tinevez et al., 2017) the foci were detected by the DoG detector, the size of the foci was set to 0.4 μm, quality threshold was set manually for each image, spanning values 2-10 depending on the image SNR. Foci were tracked using the simple LAP tracker, with the following settings that we decided upon after initial testing: linking max distance 0.4 micrometers; gap-closing max distance 0.4 micrometers; gap-closing max frame gap: 3. Filters on tracks were implemented to filter out only foci that have appeared and disappeared over the course of the time series. The tables for “tracks” were exported for each image analyzed as CSV files. The CSV files were then processed using a script written and run within the Python Google Colab Jupyter notebook environment, where the variable “track duration” was used as dwell time. To compare dynamics, dwell times were grouped into 1-s bins. Because only a small fraction of foci persisted longer than 5 s, these events were pooled into a single category. Six intervals were therefore defined: [0–1], (1–2], (2–3], (3–4], (4–5], and >5 s. To correlate foci intensity with the dwell time, the value for “mean ch1 intensity” was extracted from the “spots” CSVs. The mean intensity was then calculated for each track to which the particular foci belonged. This value was then plotted against the track’s dwell time (track duration). The dwell time of EXO84b-GFP and TPLATE-GFP foci in the ARP2/3 mutant background was analyzed using a Matlab script (Aguet et al., 2013; Johnson et al., 2020).

#### Foci density

To analyze foci density, a ROI was selected on the first focused frame of an image, and the foci were detected using the DoG detector in TrackMate (see above). The ROI area and the number of detected foci were manually recorded in a table to calculate the number of foci per 1 μm^2^.

#### Colocalization

For colocalization of foci, square ROIs were created and the foci were detected and tracked in the same way as for the dwell time analysis. Foci from both channels and for the 90° rotated control channel (negative control) were detected and tracked separately. The “spots” and CSV files were exported, and the X, Y, frame coordinates of individual foci within a track were compared using a Python script. If the foci were on the same frame with the Euclidean distance between them being less than a threshold (0.4 micrometers, which corresponds to the size of the spot), they were deemed as colocalizing. For the colocalization of foci with the cytoskeleton, square ROIs were created, the foci were detected and tracked using TrackMate as described above, and CSV files for “spots” were exported. The remaining analyses were performed using a custom Python script. Tiff images were uploaded into the environment, and the cytoskeleton channel was processed utilizing the OpenCV library as follows: the images were denoised using a median filter, contrast was enhanced using Contrast Limited Adaptive Histogram Equalization (CLAHE), and thresholded via combination of Otsu and adaptive thresholding. Based on this thresholding, binary masks of the cytoskeletal structures were generated. The X, Y, frame coordinates of individual tracked foci were compared to the cytoskeleton masks. As a negative control, the mask rotated 90° was compared as well. If the Euclidian distance between a spot and a non-zero pixel of the mask was below the threshold distance, the spot was deemed colocalizing. Further, the track was deemed as colocalizing if at least one spot of the track colocalized with the cytoskeleton. For colocalization of ER and ARP2/3, a mask of the ER was created using the trainable WEKA segmentation plugin in Fiji (Arganda-Carreras et al., 2017). TrackMate plugin (Ershov et al., 2022; Tinevez et al., 2017) was used to detect the ARP2/3 foci within (ER colocalization) or outside the ER ROI. The number of foci per area was calculated.

### Statistics and reproducibility

For the FCS data analysis, autocorrelation curves were fitted from the raw signal assuming a one-component anomalous 3D diffusion model using ZEISS ZEN software (version 3.12). A gaussian smoothing detrending filter was applied to compensate for bleaching (500 ms window). Statistical analysis of fitted data was carried out in R (version 4.5.2). Exported fitted data was standardized and quality-control filtered before statistical analysis using a threshold of 1 counts-per-molecule (CPM) as well as finite and positive diffusion coefficients. For diffusion coefficient statistical analysis, comparisons were restricted to measurements with comparable fitted molecular concentration estimate in nM. The overlap window was defined as the shared fitted concentration range between the two lines; only measurements within this range were retained for modeling (162.6-2919 nM). Candidate models were compared by maximum likelihood using AIC and likelihood-ratio tests. The additive concentration-adjusted model (logD ∼ line + log10(conc_nM) + (1 | plant_uid)) was selected, where logD is the log-transformed diffusion coefficient, line represents the induced marker (GFP vs ARPC2-GFP), log10(conc_nM) is the base-10 logarithm of the fitted FCS concentration estimate, and plant_uid a random intercept to account for biological replicate variability. Data was refitted by restricted maximum likelihood REML, and checked visually by residual diagnostics (lme4 v1.1-37, lmerTest v3.1-3, emmeans v2.0.0. performance v0.15.2). Model estimates were back-transformed for interpretation and visualized in the log scale (ggplot v4.0.0). Python scripts were used to process and analyze data for dwell times, foci densities, and colocalization. Key libraries included NumPy for numerical computations, pandas for data manipulation, SciPy and statsmodels for statistical analysis, matplotlib, and seaborn for data visualization. The methods included the Mann-Whitney U test, the Welch test, the two-sample t-test, and the paired t-test. For the dwell time data, the 95% confidence intervals of means were estimated using bootstrapping. Matlab (Aguet et al., 2013) and R (R Core Team, 2026) were used to analyze data for dwell times in ARP2/3 mutant backgrounds for EXO84b-GFP and TPLATE-GFP. Distributions of exocytic event durations were obtained for each line and compared using bootstrap resampling (1,000 iterations), with randomization based on plant or time-series identity. Confidence intervals (95%) were estimated from randomized distributions. Data were analyzed using generalized linear models (GLMs) based on the Poisson distribution. Due to overdispersion, data were log-transformed and rescaled to integers. A generalized linear mixed-effects model (GLMM) with random effects (plant, cell, series) was fitted using the glmer function from the lme4 R package. For the dwell time and foci density analyses, 3 biological replicates, 3-4 plants and 2-5 cells per plant per repetition were used for ARP2/3 subunits, NAP1-GFP, ARP2/3 subunits in mutant background, GFP-ARPC5 treated with LatB or Ory; 2 biological replicates with 2-4 plants and at least 3-5 cells per plant per repetition were used for GFP-ARPC2 in *exo84b* mutant background, GFP-ARPC5 treated with CK666, BDM, PBP. For the FRAP analysis 2 biological replicate were used with at least 2-3 plants and 2-3 cells per plant analyzed. For the colocalization of XVE::GFP-ARPC2 with mSc-ARPC5 subunits, mSc-ARPC5 with NAP1-GFP, GFP-ARPC5 with the cytoskeleton and ARP2/3 subunits with the exocyst, T-PLATE and ER-RFP markers, 2 biological replicates were used with at least 3-4 plants and 2-5 cells per plant per repetition. For the dwell time analysis of endomembrane markers in ARP2/3 mutant background, 3 biological replicates were used with at least 2 plants and 2 cells per repetition. The coimmunoprecipitation experiments were done in 3 biological replicates for the EXO84b interaction.

## Results

### The ARP2/3 complex localizes to dynamic foci in the cortical cytoplasm

In our previous work, we characterized WAVE/SCAR–ARP2/3 components associated with peroxisomes (Havelková et al., 2015; Martinek et al., 2023) and also noted a substantial cytoplasmic pool of tagged subunits of this module. In regions of the cytoplasm that appeared to contain only diffuse fluorescence by confocal microscopy, we were able to distinguish discrete submicrometer-sized foci in the cortical cytoplasm of hypocotyl and cotyledon cells using VAEM performed on TIRF microscope Zeiss Elyra (Fig. **1a**). Compared with peroxisome-associated structures, these foci were considerably smaller, were distributed homogeneously throughout the cytoplasm, and displayed a constant size of approximately 0.3 µm in diameter. Subunits of both the ARP2/3 and WAVE/SCAR complexes—specifically ARPC2, ARPC5, and native promoter-driven NAP1—formed the same cytoplasmic structures and were never observed in cells expressing GFP alone, which rules out the possibility that they represent an imaging artefact, such as nonspecific clustering of GFP-tagged proteins (Fig. **1a**). The foci exhibited high dynamics. After appearing in the cortical cytoplasm, these structures remained motionless or exhibited Brownian motion for a period of time and then disappeared (Videos S1–3). The characteristic dynamics of these structures resemble those reported for markers of endocytosis or exocytosis (Fendrych et al., 2013; Gadeyne et al., 2014; Konopka & Bednarek, 2008b, 2008a). Automated image analysis using the ImageJ TrackMate plugin (Ershov et al., 2022; Tinevez et al., 2017) allowed us to detect individual foci in single frames of a video sequence with 400 ms exposure time and track them over time. The analysis revealed that the average dwell times (the duration a particle remains in the focal plane) for *arpc5/pUBQ::GFP-ARPC5*, *arpc2*/XVE::GFP-ARPC2, and *nap1*/pNAP1::GFP-NAP1 were 1.0, 1.3, and 1.2 seconds, respectively (Table S2). The dwell times showed a quasi-exponential distribution, skewed towards the shortest ones (Fig. **1b**), similarly to markers of T-PLATE and clathrin (Aguet et al., 2013; Narasimhan et al., 2020). The domain density per μm^2^ varied across reporter lines but generally followed a normal distribution (Fig. **1c**).

**Fig. 1.**
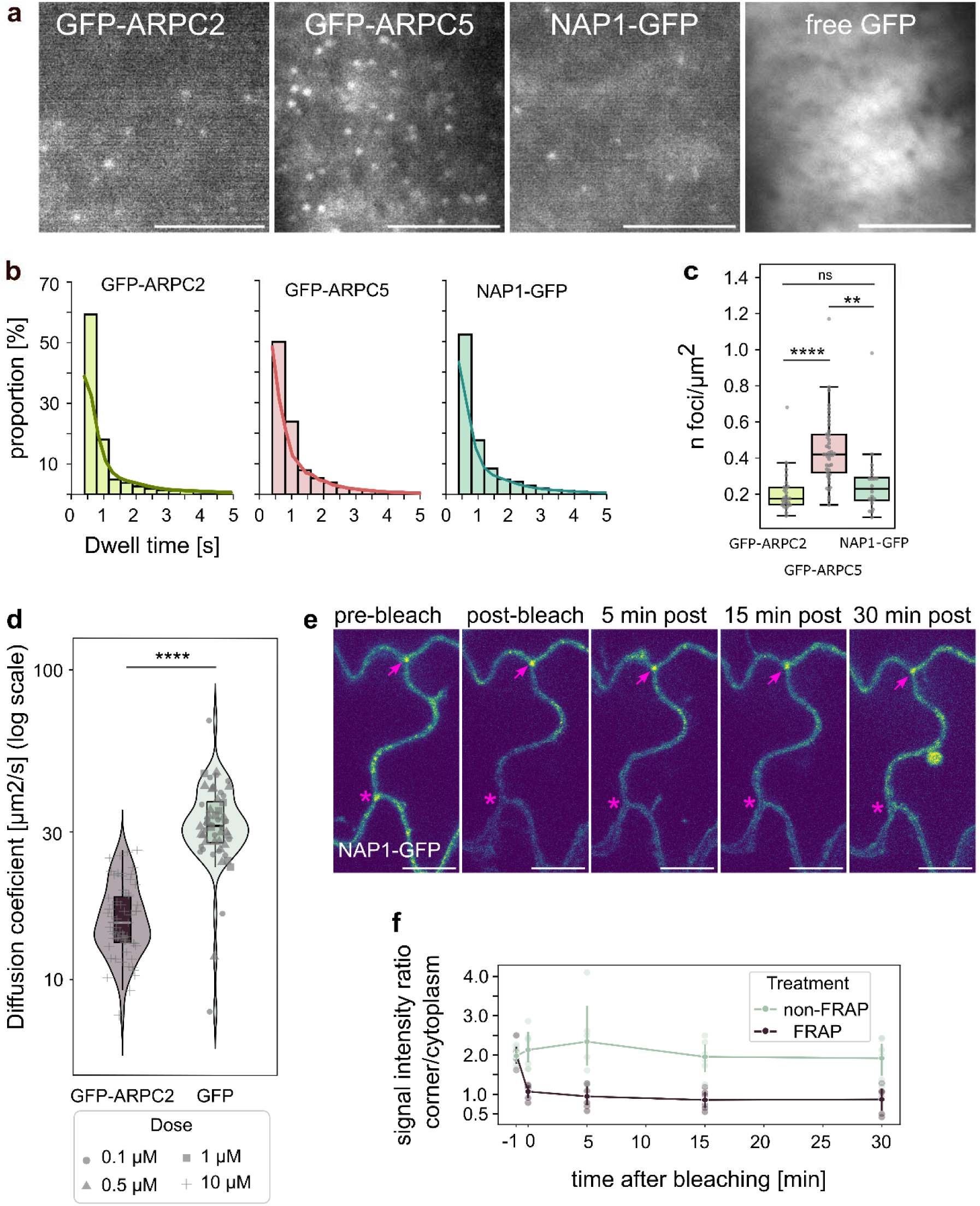
The subunits of the complexes ARP2/3 and WAVE/SCAR localize as dynamic foci in the cortical region of epidermal cell cytoplasm. (a) Visualization of *arpc2*/XVE::GFP-ARPC2, *arpc5*/pUBQ::GFP-ARPC5, *nap1*/pNAP1::NAP1-GFP and 35S::GFP using VAEM in the cortical cytoplasm of hypocotyl epidermal cells. (b) Dwell time analysis of each marker in (a) shown as dwell times distribution. For visualization purposes, only dwell times up to 5 s are shown; in rare cases foci persisted for more than 30 s (Supplemental Table S1). (c) Foci density analysis for each marker shown in (a). ARPC5 differed significantly from both ARPC2 (p < 0.001) and NAP1 (p < 0.01), whereas ARPC2 and NAP1 did not differ significantly (p > 0.05). (d) FCS analysis comparing the diffusion coefficient of *arpc2*/XVE::GFP-ARPC2 with XVE::GFP, performed at anticlinal cell walls of cotyledon epidermal cells, demonstrating reduced molecular mobility of GFP-ARPC2 (p < 0.001). Dose indicates concentration of β-estradiol used for induction. (e, f) FRAP analysis of *nap1*/pNAP1::NAP1-GFP in three-way junctions of cotyledon epidermal cells. (e) Representative image showing photo-bleached (asterisk) and control (arrow) three-way junctions. (f) Analysis of three-way junction NAP1-GFP enrichment recovery shown as signal intensity ratio of three-way junction site to the cytoplasm signal outside of the junction for photo-bleached (FRAP) and non-bleached (non-FRAP) control. While cytoplasmic signal recovered (Supplemental Figure S1H), the three-way junction enrichment did not recover within 30 minutes of analysis, indicating that NAP1-GFP localization at the three-way junction is stable rather than dynamic. Statistical analysis for plot (c) was performed using Welch’s t-test for independent samples. Statistical significance for the data in plot (d) was assessed using a concentration-adjusted linear mixed-effects model with plant identity included as a random effect (see Methods). Brightness and contrast in images (a, e) were adjusted for illustrative purposes. Scale bars: 5 µm (a) and 10 µm (e).

Hypocotyl epidermal cells were the most suitable tissue for VAEM imaging and were therefore analyzed in greatest detail. However, similar structures were also detected in epidermal cells of cotyledons (Fig. S1a). We found that the foci-to-background signal ratio was highest in cells expressing the markers at low levels (compare Fig. **1a** and Fig. S1a). Therefore, we mainly used expression constructs driven by a native promoter (*nap1*/pNAP1::GFP-NAP1), the pUBQ promoter (*arpc5*/pUBQ::GFP-ARPC5), or an inducible promoter (*arpc2*/XVE::GFP-ARPC2) stably transformed into Arabidopsis (Fig. **1a**).

The way the VAEM microscope is arranged causes only the structures closest to the cell periphery to be excited by the excitation light. As a result, the signal intensity of the fluorescent particle increases as it enters the illuminated area, thus correlating with the particle’s proximity to the PM. Aguet et al. (2013) used this phenomenon to distinguish between clathrin foci participating in active and aborted endocytic events, in which active events have longer dwell times and higher signal intensities. We investigated whether the signal intensity of ARP2/3-WAVE/SCAR markers correlated with their dwell times. We found no such correlation, indicating that the signal intensity was not dependent on the dwell time (Fig. S1b). Occasionally, we observed well-defined, high-intensity ARP2/3 foci appearing at the focal plane and then moving away from the illuminated area while losing signal (Video S2). We interpret this as the foci moving between cortical and subcortical regions.

We therefore asked whether similar structures could also be detected in other regions of the cytoplasm. However, VAEM only allows visualization of structures adjacent to the outer periclinal cell wall, and confocal laser scanning microscopy lacks sufficient spatial resolution to detect ARP2/3 foci, as mentioned above. Therefore, we employed fluorescence correlation spectroscopy (FCS) to examine the cortical cytoplasm at anticlinal cell walls. This region is of particular interest because it corresponds to the sites where adhesion defects arise in mutants of the WAVE/SCAR–ARP2/3 module. To assess the mobility of the ARP2/3 complex in this region, we analyzed the diffusion coefficient (DC) of a marker for the ARP2/3 subunit ARPC2 and compared it with the DC of free GFP. Because ARP2/3 functions as part of a cytoplasmic protein complex, it is expected to exhibit different mobility than freely diffusing GFP. Both markers were expressed under inducible promoters (XVE::GFP-ARPC2 and XVE::GFP), which allowed us to control expression levels and maintain comparable cytoplasmic concentrations of the two markers, thereby minimizing concentration-dependent artifacts. FCS measurements revealed that the diffusion coefficient of GFP-ARPC2 was substantially lower than that of free GFP (Fig. **1d**; Fig. S1c), indicating reduced molecular mobility and suggesting that the protein diffuses as part of a larger molecular complex. These results strongly suggest that the foci observed in the cortical cytoplasm adjacent to the outer periclinal cell wall were also present in other regions of the cytoplasm.

WAVE/SCAR accumulates at the three-way cell junctions of epidermal cells (Dyachok et al., 2008; Qin et al., 2021; Wang et al., 2016) (Fig. S1e; Video S4). This pattern is particularly important, as these domains correspond to regions where adhesion defects are observed in mutants lacking functional ARP2/3. Importantly, three-way junction localized WAVE/SCAR represent functional and active domain of WAVE/SCAR, because the loss of the BRK1 subunit causes mislocalization of NAP1 (Fig. S1e, f; Videos S4, 5). We investigated whether the formation of these accumulations requires the high dynamics of the complex that we observed in the cortical cytoplasmic layer. Using the *nap1*/pNAP1::NAP-GFP line, we performed fluorescence recovery after photobleaching (FRAP) analysis at three-way junctions. During 30 minutes of recovery after photobleaching, no restoration of accumulation at the three-way junctions was observed, whereas diffuse cytoplasmic signal clearly recovered (Fig. **1e, f**; Fig. S1d). This result shows that the three-way cell junction localization must represent a more stable fraction of the NAP1 subunit pool, possibly bound to the PM, in contrast to the extremely dynamic structures observed in VAEM (Fig. **1a, b**; Video S3).

### Cortical cytoplasm contains both assembled and partially assembled ARP2/3 complexes

The ARP2/3 complex consists of seven subunits, all except ARPC3 being required for its function in plants. To gain an insight whether the assembly status of the complex affects the foci, we expressed GFP-ARPC5 or GFP-ARPC2 subunits in *arpc2* and *arpc5* mutants, respectively, to produce non-rescued plants with an ARP2/3 marker. We analyzed the cortical foci in these lines and compared the results with the rescued *arpc2*/XVE::GFP-ARPC2 and *arpc5*/pUBQ::GFP-ARPC5. Surprisingly, some dynamic foci were also detected in the cortical cytoplasm of the non-rescued mutants (Fig. **2a** and Fig. S2a). However, analysis of dwell times revealed a change in dynamics - the proportion of foci with the shortest dwell times increased in the non-rescued mutant background compared with the corresponding rescued lines (Fig. **2b**; Fig. S2b).

**Fig 2.**
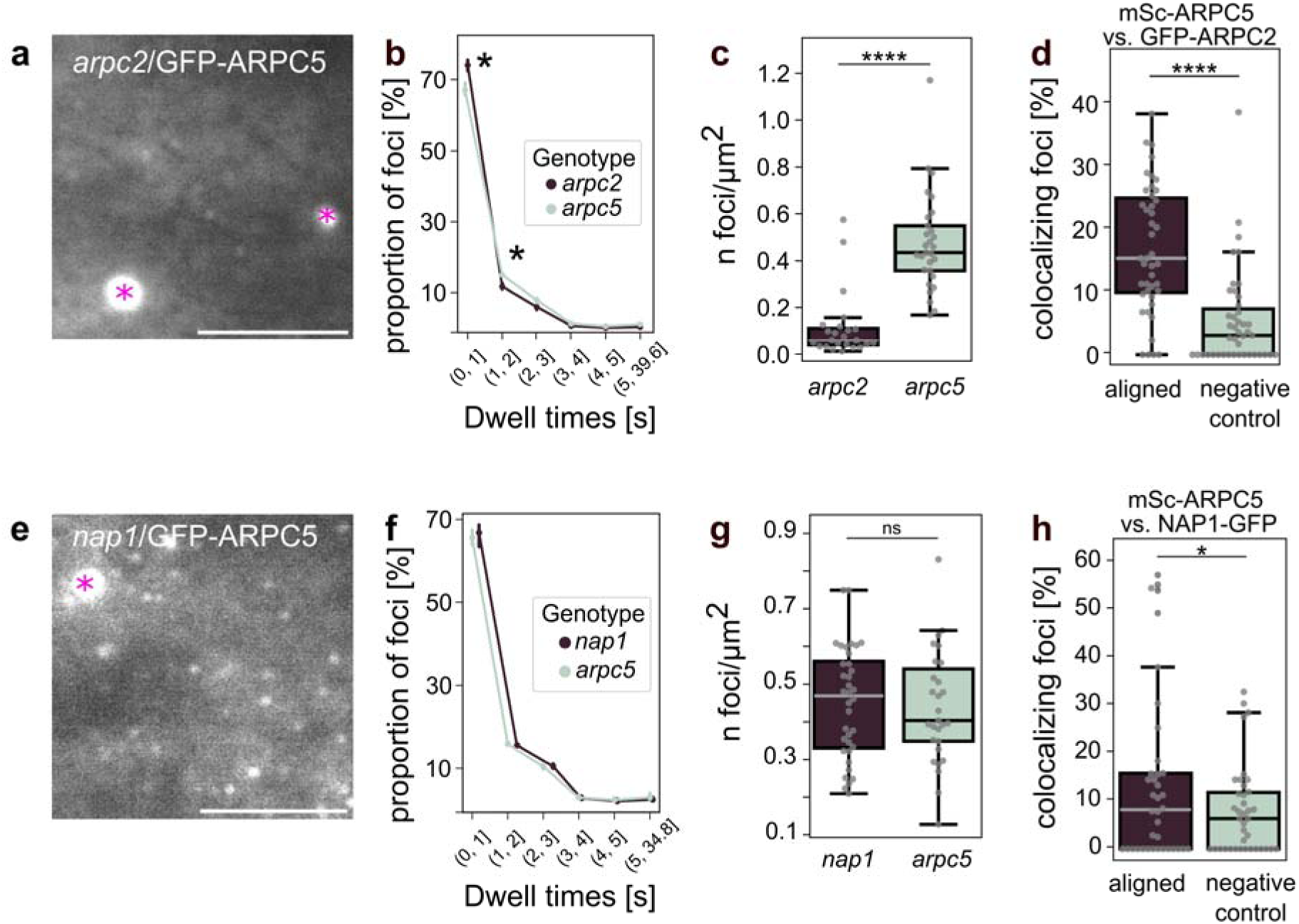
The amount and dynamics of ARP2/3 is dependent on its assembly status but not its activation. (a) *arpc2*/pUBQ::GFP-ARPC5 in the cortical cytoplasm (VAEM, hypocotyl pavement cell). (b) Analysis of pUBQ::GFP-ARPC5 foci dwell time in rescued (*arpc5*) and non-rescued (*arpc2*) mutant background, shown as a proportion of foci with certain dwell time (binned). Asterisks indicate statistically significant differences in foci proportions (bootstrapped confidence intervals are not overlapping). (c) The density of pUBQ::GFP-ARPC5 foci in *arpc2* mutant is lower than in rescued *arpc5* mutant (p < 0.0001). (d) The colocalization analysis of 35S::mSc-ARPC5 and XVE::GFP-ARPC2 foci, shown as the percentage ARPC5 foci colocalizing with ARPC2 foci compared to negative controls (90 ° rotated image; p < 0.0001). (e) *nap1*/pUBQ::GFP-ARPC5 in the cortical cytoplasm (VAEM, hypocotyl pavement cell). (f) Analysis of pUBQ::GFP-ARPC5 foci dwell time in rescued (*arpc5*) and non-rescued (*nap1*) mutant background, shown as a proportion of foci with certain dwell time (binned); the difference is not significant (in all cases the bootstrapped confidence interval are overlapping). (g) The density of pUBQ::GFP-ARPC5 foci in *nap1* mutant and rescued *arpc5* mutant is comparable (p-value > 0.05). (h) The colocalization analysis of 35S::mSc-ARPC5 with pNAP1::NAP1-GFP, shown as the percentage ARPC5 foci colocalizing with NAP1 foci compared to negative controls (90 ° rotated image; p < 0.05). Statistical analysis: (b, f) bootstrapping with error bars around timepoint values representing the 95% confidence intervals; (c, g) Welch’s t-test for independent samples; (d, h) Wilcoxon test. (a, e) Scale bar: 5 mm; asterisks indicate peroxisome-associated ARPC5, excluded from the analysis; brightness and contrast have been modified for illustrative purposes.

Although the difference was statistically significant only for the ARPC5 subunit, the ARPC2 followed a similar trend (Fig. **2b** and Fig. S2b). Importantly, in non-rescued lines, the density of foci decreased significantly compared to the respective rescued ones (Fig. **2c** and Fig. S2c). We also observed a decrease in the signal intensity of the foci relative to the background in non-rescued lines (Fig. S2g), similarly to 35S-driven ARP2/3 markers (Fig. S1a). This could indicate an excess of free subunits due to their inability to assemble into the complex. These results demonstrate that some of the observed foci, even in a rescue background, may represent incomplete ARP2/3 complexes. To further investigate whether the observed foci represent fully assembled ARP2/3 complexes or only partially assembled complexes, we co-expressed the markers XVE::GFP-ARPC2 and 35S::mSc-ARPC5 and analyzed their colocalization.

The analysis was performed using a modified approach in which foci were tracked separately in both channels using TrackMate, followed by colocalization analysis based on their x, y, and t coordinates using a custom Python script. The colocalization values obtained from the aligned channels were compared with those acquired from the same time sequences in which one channel was rotated by 90°, serving as a negative control. This analysis demonstrated that, on average, 15 % of ARPC5 foci co-localized with the ARPC2 foci, while only on average 4 % colocalized in our rotated negative control (Table S3; Fig. **2d**; Fig. S2d, e). While the colocalization events observed in the negative control samples result from random overlap of structures caused by their high density in both channels, the statistically significant colocalization between the two ARP2/3 markers confirms that the two subunits physically co-localize in cortical dynamic foci. Surprisingly, however, the analysis also showed that the majority of foci did not contain both markers, further suggesting the presence of a large pool of partially or incompletely assembled ARP2/3 complexes in the cytoplasm of plant cells.

### A small fraction of cortical ARP2/3 foci represent potentially active complexes

The dynamics of the functional interaction between these two complexes is studied mainly using *in vitro* (Smith, Daugherty-Clarke, et al., 2013; Smith, Padrick, et al., 2013) and *in silico* approches (Iyer et al., 2025). It is clear that to nucleate branched actin filament, the assembled ARP2/3 complex must be activated by the WAVE/SCAR activating complex. The activating complex however dissociates from ARP2/3 upon its binding to the mother actin filament and the start of new filament nucleation (Iyer et al., 2025; Smith, Padrick, et al., 2013). Thus, ARP2/3 complexes not associated with WAVE/SCAR may represent either inactive complexes or complexes that have already initiated actin nucleation, whereas ARP2/3 complexes associated with WAVE/SCAR represent nucleation-competent complexes. We asked whether markers for ARP2/3 and WAVE/SCAR colocalize within the detected cytoplasmic structures and whether the cortical foci can also be detected in mutants lacking a functional WAVE/SCAR complex. To this end, we expressed ARP2/3 marker pUBQ::GFP-ARPC5 in the *nap1* mutant background (Fig. **2e**). The analysis of the dynamics and density of ARPC5 foci revealed that these values are comparable for *nap1* mutant and rescued *arpc5* line (Fig. **2f, g**). This indicates that ARP2/3 complex and subcomplex formation is independent of the activation status. To visualize possible ARP2/3 and WAVE/SCAR complexes interaction, we co-expressed ARP2/3 marker mSc-ARPC5 and WAVE/SCAR marker NAP1-GFP and analyzed their colocalization in the cortical region. Indeed, the two markers co-localized in dynamic foci, though to a lesser extent than that of the two ARP2/3 subunits; on average, 14,03% of mSc-ARPC5 foci colocalized with NAP1-GFP (Table S3; Fig. **2h**; Fig. S2d, f). The cortical ARP2/3 foci thus possibly form active and inactive pools. These results indicate that the majority of ARP2/3 foci observed in plants with a fully functioning complex might represent incomplete and likely inactive assemblies. Also, only a small fraction corresponds to the ARP2/3 complex bound to the activation complex representing potentially active complexes.

### Dynamic ARP2/3 foci interact with the cortical cytoskeleton

The above described observations suggested that only a small fraction of the ARP2/3 complexes observed in the cortical cytoplasm are likely to represent active complexes. Because ARP2/3 complex function is closely linked to the actin cytoskeleton, our subsequent analysis focused on whether ARP2/3 foci colocalize with the cortical actin filaments (AFs). In earlier studies, we also demonstrated that plant ARP2/3 specifically colocalizes with cortical microtubules (MTs) *in vivo* (Havelková et al., 2015); therefore, we also analyzed the relationship between dynamic cortical ARP2/3-containing foci and microtubules.

Colocalizing the different structures *in vivo* was a challenging task because both the foci and cytoskeleton are very dynamic (Videos S6, 7). We used the pUBQ::GFP-ARPC5 marker, which provided the best signal-to-noise ratio for colocalization with Lifeact-RFP and mCh-TUA5 (Fig. **3a**). Colocalization was assessed using the spot tracking approach described above, combined with the matching of the x, y, t coordinates of individual foci to binary masks of the cytoskeletal channel (Fig. S3a, b). As in previous analyses, videos with one channel rotated by 90° served as a negative control. Using this approach, we were able to demonstrate that ARPC5 colocalized with both AFs and MTs. The mean colocalization rate was 46,32 % in aligned images and 37,93 % in the negative control for AFs (Fig. **3b**; Table S4), and 73,01 % in the aligned and 64,12 % in the negative control for MTs (Fig. **3c**, Table S4). Analogically to foci colocalization analysis described above, observed cytoskeleton-localized foci represented were a mixture of random and true colocalizations due to the high density of both structures. The true colocalizations can be very roughly estimated by subtracting the negative control value from the experiment, which is 8,4 % ARPC5 foci colocalized with AFs and 8,97% with MTs. These values are, of course, only estimates; nevertheless, they suggest that cortical ARP2/3 foci have a spatial association with the cytoskeleton.

**Fig. 3.**
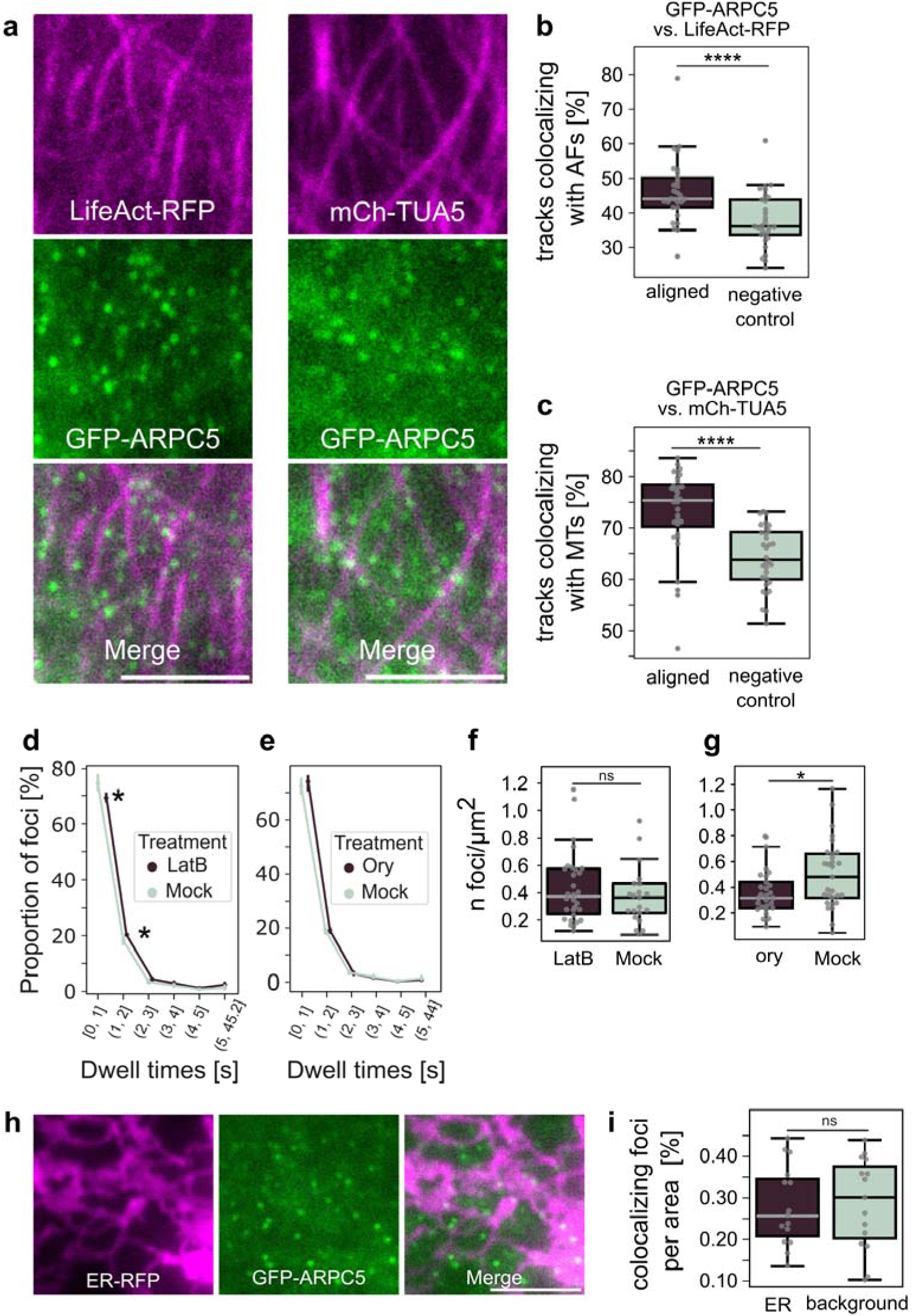
Cytoskeleton-dependent ARP2/3 localization and dynamics. (a) 35S::LifeAct-RFP for actin filaments (AF) visualization and pUBQ::GFP-ARPC5 and 35S::mCh-TUA6 for microtubules (MTs) visualization and pUBQ::GFP-ARPC5 colocalization in epidermal hypocotyl cell (VAEM). (b) Analysis of GFP-ARPC5 colocalization with AFs shown as a percentage of GFP-ARPC5 foci colocalizing with actin filaments (p < 0.0001). (c) Analysis of GFP-ARPC5 colocalization with MTs shown as a percentage of GFP-ARPC5 foci colocalizing with MTs filaments (p < 0.0001). (d, e) Analysis of dynamics of pUBQ::GFP-ARPC5 in latrunculin B (LatB)-treated plants (D; 2h, 2 µM) and oryzalin (Ory)-treated plants (E; 2h, 20 µM) shown as a proportion of GFP-ARPC5 foci with certain dwell time (binned). LatB treatment resulted in the reduced amount of short dwell time-foci, while oryzalin had no effect on ARPC5 dynamics (asterisks label statistically significant instances where bootstrapped confidence intervals are not overlapping). (f, g) Analysis of pUBQ::GFP-ARPC5 foci density in LatB-treated plants (F; 2h, 2 µM) and oryzalin-treated plants (g; 2h, 20 uM). While LatB had no effect on foci density (p > 0.05), oryzalin promoted their amount (p < 0.05). (h, i) Analysis of cortical ER and ARP2/3 cortical foci with ER. (h) Colocalization of ER-RFP and pUBQ::GFP-ARPC5 (cotyledon pavement cell). (i) Analysis of GFP-ARPC5 colocalization with ER shown as percentage of ARPC5 foci colocalizing either with the ER marker or background demonstrated no significant colocalization with ER membranes (p > 0.05). Statistical analysis: a paired t-test (b, c); bootstrapping - lines around timepoint values represent the 95% confidence intervals (if not overlapping, the result is considered statistically significant) (d, e); Mann-Whitney-Wilcoxon test (f, g); Welch’s t-test for independent samples (i). Brightness and contrast have been modified in images (a, h) for illustrative purposes. Scale bar: 5 µm.

Functional interactions between ARP2/3 foci and the cytoskeleton were tested under conditions of cytoskeleton depolymerization. We analyzed the number and dynamics of cortical ARP2/3 foci in the pUBQ::GFP-ARPC5 marker line following treatment with the cytoskeleton-disrupting drugs latrunculin B (LatB; Morton et al., 2000) and oryzalin (Ory; Hugdahl & Morejohn, 1993). Analysis of dynamic properties showed that LatB-treated cells exhibited a reduced proportion of foci with the shortest dwell times (Fig. **3d**), suggesting partial stabilization of foci in the absence of actin filaments. In some cases, foci in LatB-treated plants appeared arranged in patterns reminiscent of microtubules (Fig. S3c). However, the total number of foci remained unchanged in LatB-treated cells (Fig. **3f**). Interestingly, ARP2/3 foci in oryzalin-treated plants showed unchanged dynamics (Fig. **3e**), but their abundance was significantly reduced compared to untreated controls (Fig. **3g**). Further we tested the effect of a specific ARP2/3 complex inhibitor CK666 (Nolen et al., 2009) and two myosin inhibitors, BDM and PBP (Siegman et al., 1994; W. Zhang et al., 2019), on the dynamics and density of ARP2/3 structures in the cortical cytoplasm. Despite the reported effects of CK666 and PBP on actin dynamics (Xu et al., 2024), we did not observe any changes in either foci dwell time or density in plants treated with 10 µM CK666 (Fig. S4a-c). Similarly, treatment with either 10 µM PBP or 50 mM BDM had no effect (Fig. S4d-i).

### Cortical ARP2/3 foci have a strong link to exocytosis

Visualization of the ARP2/3 complex in several studies has revealed a functional link between ARP2/3, PM and the cortical ER. Wang et al. (2016, 2019) identified cortical ARP2/3-containing structures together with proteins involved in the formation of ER–PM contact sites (EPCS), such as VAP27, AtEH/Pan1, and actin, and proposed that ARP2/3 may specifically recruit actin to these sites to connect endocytosis and autophagy (Wang et al., 2016). Therefore, we analyzed the colocalization between the dynamic foci detected by VAEM in our study and the ER. Our analysis showed that the occurrence of ARP2/3 foci in the cortical cytoplasm is independent of ER membranes, as the density of cortical ARP2/3 foci was similar in regions with and without ER membranes (Fig. 3**h, i**). The homogeneous distribution of cortical ARP2/3 and the lack of its specific accumulation at ER membranes suggest that the observed structures are not part of EPCS, but rather represent independent structures.

In subsequent experiments, we analyzed the relationship between cortical ARP2/3 foci and the processes of endocytosis and exocytosis. We focused on these processes for several reasons. First, the characteristic pattern of these foci and their dynamics— namely their transient localization at the cell periphery and minimal lateral movement— closely resemble the behaviour of markers of endocytosis or exocytosis at the cell cortex (Aguet et al., 2013; Fendrych et al., 2013; Konopka & Bednarek, 2008b; Narasimhan et al., 2020). Furthermore, components of the endocytic machinery have been shown to colocalize with and be functionally linked to the ARP2/3 complex in a previous study (Wang et al., 2019). Another reason for our interest in these processes is the fact that these membrane-associated events could, in principle, involve the actin meshwork in the cortical cytoplasm, such as Arp2/3-actin function in endocytic processes in yeast (Young et al., 2004). To explore the relation of ARP2/3 to these processes in the cortical cytoplasm, we co-expressed markers for endocytosis TPLATE-RFP and DRP1C-mOra and for exocytosis EXO84b-RFP with GFP-tagged subunits of ARP2/3 complex (Fig. **4a**, Fig. S5e). Using the colocalization approach described above including comparison with negative controls we found significant colocalization for all tested markers (Fig. **4b, c**; Fig. S5c, d, f; Table S5). We found that the colocalization rate was consistently higher for the exocytosis marker than for the endocytosis markers. After establishing that markers of endocytosis and exocytosis exhibit a significant degree of colocalization with ARP2/3 foci, we asked whether these processes are functionally dependent on ARP2/3-mediated actin polymerization. We expressed the markers TPLATE–GFP and EXO84b–GFP in *arpc5* mutants and analyzed the dynamics of these processes by measuring the dwell time of structures corresponding to endocytic and exocytic events in the cortical cytoplasm, visualized by spinning disc.

**Fig. 4.**
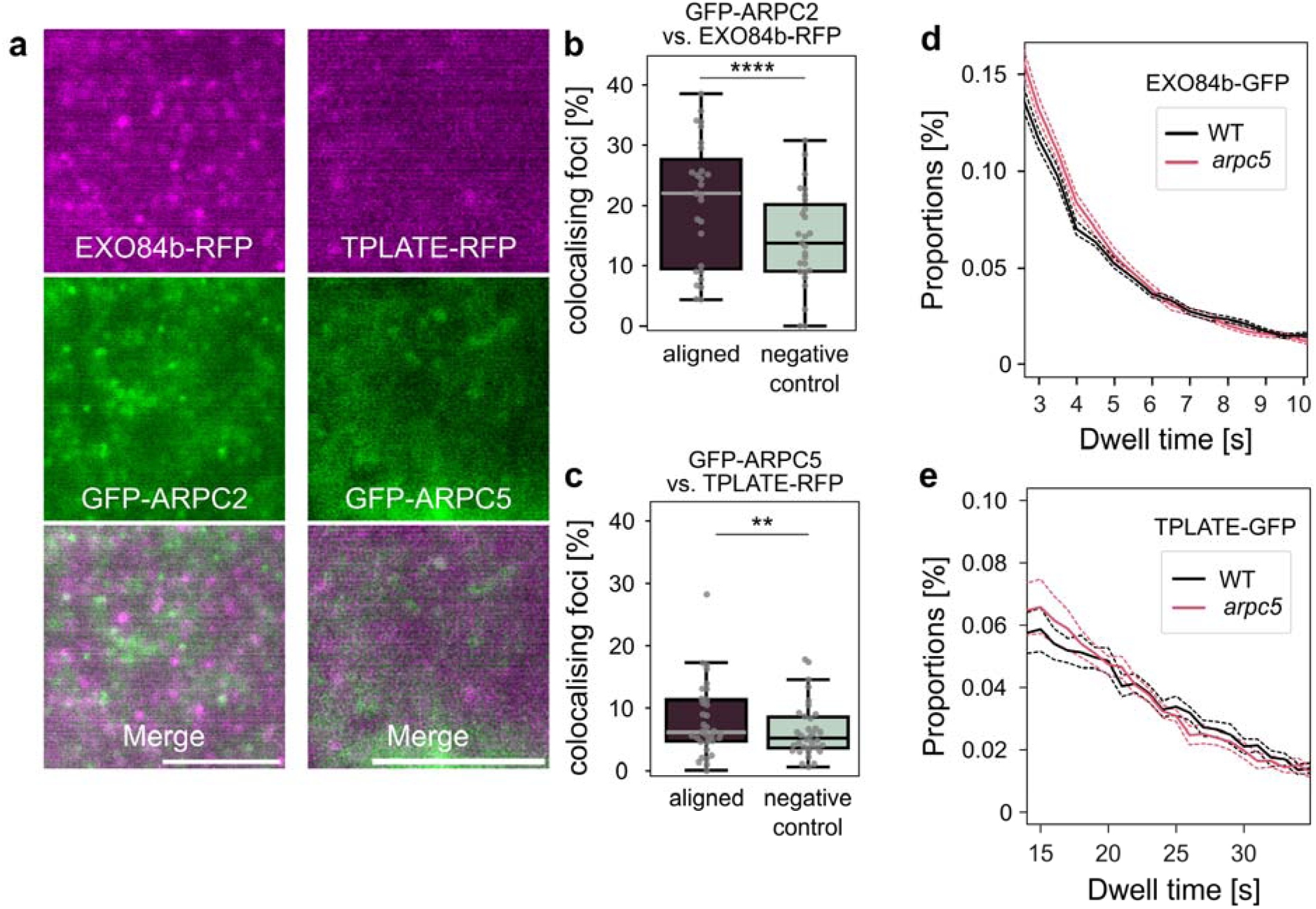
Cortical ARP2/3 colocalizes with membrane trafficking markers and is functionally linked to exocytosis. (a) 35S::EXO84b-RFP and XVE::GFP-ARPC2 and 35S::TPLATE-RFP and 35S::GFP-ARPC5 colocalization in epidermal hypocotyl cell (VAEM). (b, c) Analysis of ARP2/3 and membrane trafficking markers colocalization shown as a percentage of GFP-ARPC2 foci colocalizing with EXO84b-RFP compared to negative controls (90 ° rotated image; p < 0.0001) (b) and GFP-ARPC5 foci colocalizing with TPLATE-RFP compared to negative controls (90 ° rotated image; p < 0.01) (c) demonstrated significant colocalization of ARP2/3 and membrane trafficking markers. (d, e) Dwell time analysis of exocytosis GFP-EXO84b (d) and endocytosis TPLATE-GFP (e) foci at the plasma membrane shown and a proportion of foci with certain dwell times. While the amount of short GFP-EXO84b foci dwell times significantly increased in *arpc5*, compared to wt (d), the proportion of TPLATE-GFP foci with certain dwell times in wt and *arpc5* was comparable (e). Statistical analysis: Wilcoxon test for paired samples (b, c), bootstrapping - dashed lines around the solid ones represent the 95% confidence intervals (if not overlapping, the result is considered statistically significant) (d, e). Brightness and contrast have been modified for images (a) for representative reasons. Scale bar: 5 µm.

By comparing the dynamics of exocytic events (EXO84b–GFP) in the cortical cytoplasm between wild-type plants and *arpc5* mutant, we found that the number of events with very short dwell times was higher in the mutant lacking a functional ARP2/3 complex (Fig. **4d**). In contrast, comparison of the dynamics of the endocytic marker (TPLATE– GFP) between wild-type and *arpc5* mutant background did not reveal a significant difference in the distribution of dwell times (Fig. **4e**).

Encouraged by results suggesting a functional link between the dynamics of exocytosis markers and ARP2/3 activity, we investigated whether the occurrence and dynamics of cortical ARP2/3 foci are affected by a non-functional exocyst complex in the *exo84b* mutant. To this end, we expressed the ARP2/3 subunit XVE::ARPC2–GFP in the *exo84b* mutant background and compared its dynamics between wild-type and mutant lines. Neither the density nor the dynamics of ARPC2–GFP in the *exo84b* mutant background were altered, suggesting that ARP2/3 function is upstream to EXO84b (Fig. S5a, b). We next sought to determine whether the functional interaction between the exocyst subunit EXO84b and the ARP2/3 complex also involves a direct physical association between these components. Indeed, biochemical analysis demonstrated that these proteins interact directly *in vivo*. Pull-down assays from extracts of plants co-expressing EXO84b–RFP and the ARP2/3 subunits ARPC2 or ARPC5 revealed that both subunits co-immunoprecipitated with EXO84b (Fig. **5**).

**Fig. 5.**
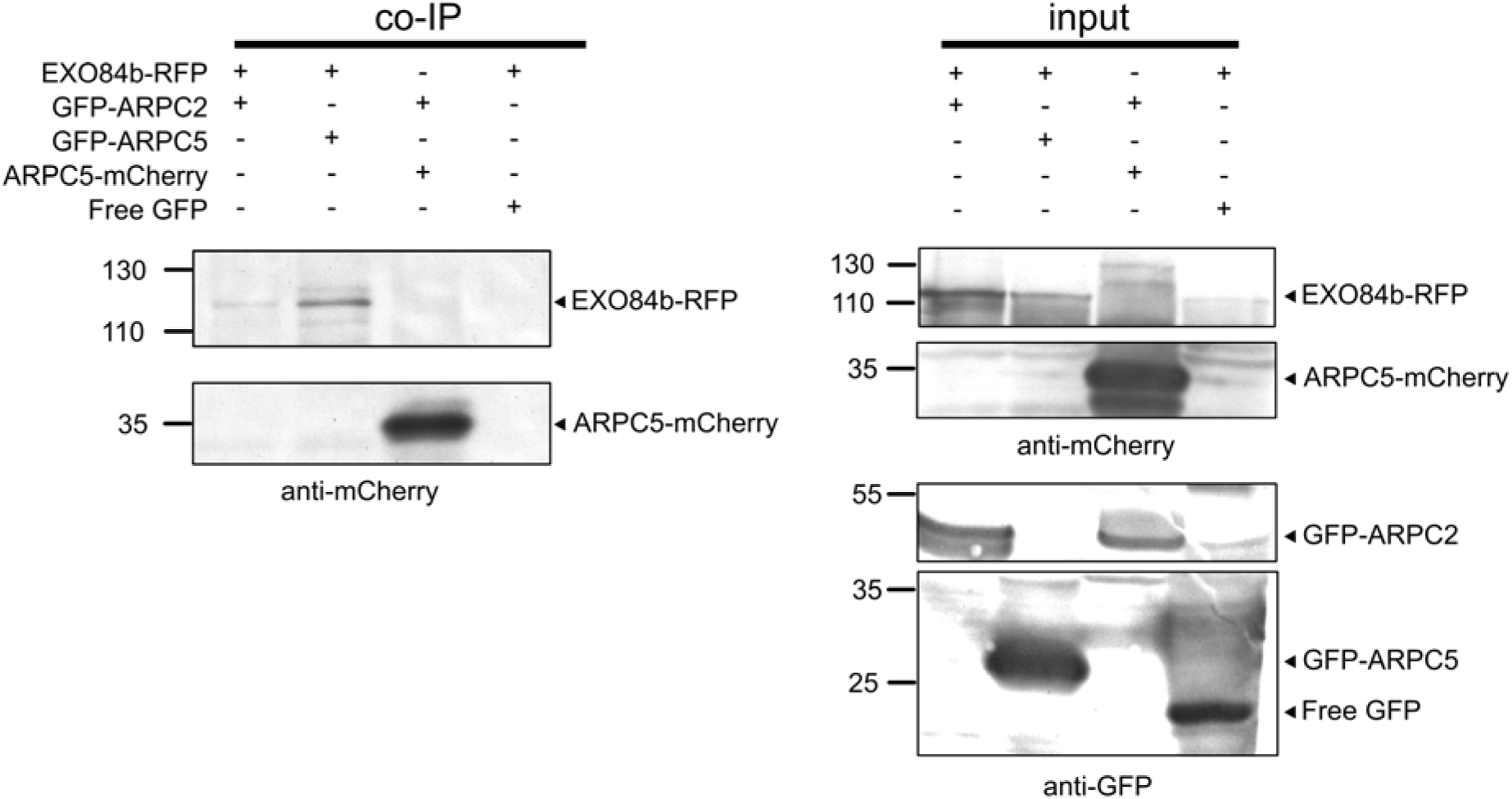
Biochemical analysis of EXO84b interaction with ARP2/3. Co-immnoprecipitation assay of protein extracts isolated from Arabidopsis plants co-expressing EXO84b-RFP with GFP-ARPC2, GFP-ARPC5 or GFP, and plants co-expressing GFP-ARPC2 and ARPC5-mCherry as a positive control. Extracts were mixed with magnetic beads coated with anti-GFP antibody and co-immunoprecipitated proteins were probed using anti-mCherry antibody (left panel, co-IP). Input protein extracts were probed using anti-mCherry and anti-GFP antibody (right panel, input). Blots were cropped; original uncropped blots are shown in Figure S6a.

## Discussion

### Subunits of WAVE/SCAR-ARP2/3 module form dynamic cortical foci

Only few works have quantitatively examined the dynamics of ARP2/3-associated structures near or at the plant plasma membrane. Previous *in vivo* observations reported pressure-induced puncta in epidermal cells involved in autophagy (Wang et al., 2016, 2019). Tips of growing trichomes contain dynamic ARP2/3 foci (Yanagisawa et al., 2018). WAVE/SCAR-ARP2/3 is associated with motile peroxisomes as puncta of varying sizes (Martinek et al., 2023). Our current work provides evidence that within the cytoplasmic membrane and proximal cortical cytoplasm, components of the plant WAVE/SCAR–ARP2/3 machinery exists as an additional population of abundant, highly dynamic, short-lived foci. Their dynamics and remarkably uniform size suggest that these structures represent either individual ARP2/3 complexes or discrete foci containing a defined number of complexes. Their distinct nature is further brought out by the fact that they are not resolved by standard confocal microscopy, likely due to their small size and rapid dynamics. The quality of the observations depended on the promoter driving marker expression. Markers whose expression was driven by native, UBQ, or inducible XVE promoters gave much better resolution than those driven by the 35S promoter due to limited diffuse background fluorescence and increased resolution. Although the cortical foci are at the edge of detectability, they are unlikely to represent imaging artefacts. This is supported by the observation that WAVE/SCAR foci are detectable in the *nap1*/pNAP1::NAP1-GFP line (Wang et al., 2016), whereas such cortical foci were never observed in cells expressing free GFP. Further, FCS analysis suggests ARPC2 clustering in other parts of the cytoplasm, inaccessible to VAEM, which further supports their native character. Moreover, foci abundance was changed in mutants lacking an active ARP2/3 complex, and we could modulate their abundance and dynamics by application of anti-cytoskeletal drugs.

Based on their size and dynamics, the structures observed in the cortical cytoplasm in VAEM most closely resemble the dynamic foci found at the apex of growing trichomes (Yanagisawa et al., 2018). In contrast, the structures observed in other studies in epidermal cells (Martinek et al., 2023; Wang et al., 2016, 2019) do not match the properties of these dynamic foci, as they are substantially larger, readily detectable by confocal microscopy, and inducible by mechanical stress (Wang et al., 2016, 2019) or associated with peroxisomes (Martinek et al., 2023). As the distribution of the cortical cytoplasmic foci does not depend on the ER, we currently lack evidence that these foci correspond to EPCS (Wang et al., 2016, 2019).

### Most ARP2/3 foci likely represent partially assembled and non-activated subcomplexes

Visualization of a relatively large area of the cortical cytoplasm in epidermal cells by VAEM imaging enabled automated particle tracking of these fast cortical events, yielding robust quantitative data that formed the basis for the characterization of these dynamic protein assemblies at the plant cell surface. The analysis of different subunits co-expression in wild-type and mutant backgrounds showed that only a minority of foci contain assembled ARP2/3 or the activator subunits indicating that plants maintain a large pool of pre-assembled inactive ARP2/3 complexes. This view is broadly consistent with earlier studies but substantially extends them through quantitative *in vivo* analysis. Immunolocalizations have already revealed a pattern of puncta of varying sizes, suggesting that ARP2/3 and WAVE/SCAR exist in excess (C. Zhang, Mallery, & Szymanski, 2013; C. Zhang, Mallery, Reagan, et al., 2013). Our quantitative *in vivo* analysis of ARP2/3 confirmed the findings of biochemical studies that cytoplasmic plant ARP2/3 may consist of a mixture of assembled and partially assembled populations and that the assembly of ARP2/3 complex is independent of its activation status (Kotchoni et al., 2009), as the ARP2/3 foci density dramatically lowered in both *arpc2* and *arpc5* mutants, but not in *nap1*. A similar effect of a single *exo70A1* subunit mutation on the occurrence of cortical foci was recently described for the exocyst complex (Fendrych et al., 2013; Jiang et al., 2025; Synek et al., 2021). The small subpopulation of cortical ARP2/3 observed to interact with WAVE/SCAR likely represents the nucleation-competent complex, while ARP2/3 without interaction can be either inactive or currently nucleating ARP2/3 complex already dissociated from WAVE/SCAR (Iyer et al., 2025; Smith, Padrick, et al., 2013).

The high abundance of ARP2/3 and WAVE/SCAR foci however poses a limitation for the quantification of microscopy data. The high particle density inevitably inflates apparent co-localization frequencies. Although rotated-channel negative controls mitigate this effect, higher-resolution approaches will be required to resolve stoichiometry within individual foci.

### Both actin and microtubule cytoskeletons colocalize with cortical ARP2/3 foci

Extending this dynamic view, our cytoskeletal colocalization experiments further elucidate the ARP2/3-cytoskeletal functional context. Only a subset of foci overlapped with detectable actin, supporting the view that most assemblies are not actively nucleating filaments, but are rather part of a pre-assembled pool. The effect of ARP2/3 absence on the branching of actin filaments within the cortical region has been clearly demonstrated (Xu et al., 2024). However, we could not establish a strong link between cortical foci and active actin branching due to the high dynamics of both structures. Perhaps simultaneous visualization of ARP2/3, WAVE/SCAR and the cytoskeleton could help distinguish the pre-activated complexes and complexes participating in actin nucleation. As the *in vitro* and *in silico* molecular dynamics assays suggest, ARP2/3 activation by NPF is a multistep process, during which both ARP2/3 and NPF transition through several conformation states. The ARP2/3 is bound to the WAVE/SCAR only transiently, as the activation complex rapidly dissociates before the daughter actin filament elongation commences (Iyer et al., 2025; Smith, Padrick, et al., 2013). Whether it is possible to use *in vivo* imaging to discern the individual states, including the one where ARP2/3 is currently nucleating an actin filament remains a question.

Also, the current toolkit of actin reporters in plants remains insufficient to unambiguously capture very transient nucleation events at the cortex. Based on our data, we conclude that a relatively small but significant fraction of cortical ARP2/3 complexes colocalizes with actin *in vivo*, and that both systems undergo extremely rapid remodeling in the cytoplasm, which limits the resolution of potential nucleation events. The association with MTs is a significant recurring observation (Havelková et al., 2015; C. Zhang, Mallery, & Szymanski, 2013), and further supports the possibility of indirect coordination between the two cytoskeletal systems. Drug treatments showcased the differential roles of the cytoskeleton: actin depolymerization primarily altered foci dynamics, whereas microtubule depolymerization reduced their density. This finding is particularly intriguing in light of the colocalization of the WAVE/SCAR–ARP2/3 module with the exocytic marker EXO84b. Cortical MTs define microtubule-determined zones in which exocytosis preferentially occurs (Lindeboom et al., 2026). Because these zones extend beyond the width of individual MTs, preferential exocytosis is unlikely to result simply from direct vesicle association with MTs, and the mechanism maintaining the identity of these zones remains unknown. Coordination of exocytosis, actin, and MTs through ARP2/3-mediated actin nucleation could contribute to this mechanism.

### Cell edges are sites of stable localization of the WAVE/SCAR complex subunit NAP1

Using FRAP approach, we demonstrated that NAP1 accumulation at three-way junctions (Chi & Ambrose, 2025; Dyachok et al., 2008; Wang et al., 2016) represents a highly stable localization of the WAVE/SCAR complex. Based on the fact that unlike WAVE/SCAR, ARP2/3 markers do not accumulate at three-way junctions, we hypothesize that local ARP2/3 activation at these sites is transient and is spatially defined by a more stable and plasma membrane-localized WAVE/SCAR. Unfortunately, anticlinal cell walls are beyond the reach of VAEM, which would be probably effective in detection of ARP2/3 dynamic population.

### Cortical dynamic WAVE/SCAR–ARP2/3 foci associate with endocytic and exocytic markers

Stable character of NAP1-GFP at three-way cell junctions and the dynamic foci in the cortical cytoplasm indicate distinct functional roles of these two types of assemblies. The stochastic dynamics of the cortical WAVE/SCAR–ARP2/3 module are consistent with the behaviour of the actin cytoskeleton itself in the cortical cytoplasm (Staiger et al., 2009), and even a smaller subpopulation of active complexes may have a specific functional role. We identified such a specific role in our study, as the abundant, short-lived assemblies colocalized with endocytic and exocytic markers. The observed colocalization with an exocytic marker has a functional basis, as we demonstrated statistically significant modulation of exocytosis in mutants lacking the ARP2/3 complex, whereas endocytic processes were unaffected in our experiments. The interaction between ARP2/3 and the exocyst is not unique to plants, as Exo70, a subunit of the exocyst complex, interacts with Arp2/3 in mammalian cells (X. Zuo et al., 2006) and stimulates its activity by promoting the interaction of Arp2/3 with its NPF WAVE2 (Liu et al., 2012). The spatial proximity of a subset of foci to an exocytic marker at the periphery of plant cells points to a potential role in secretion-associated membrane remodeling. The exocyst is a key tethering complex required for polarized exocytosis and cell surface expansion, and its functional interplay with the actin cytoskeleton has been implicated in plant cell wall synthesis (Hála et al., 2008; Jiang et al., 2025; Žárský et al., 2013; W. Zhang et al., 2019, 2021). The significance of the exocyst-actin interaction is supported by observed association of exocyst with myosin motor XIK at sites of vesicle exocytosis at the PM (W. Zhang et al., 2021) as well as by the effect of LatB-mediated actin disruption on both the localization and dynamics of EXO84b cortical foci (Fendrych et al., 2013). Notably, the dwell times of EXO84b events measured in our experiments were, on average, shorter and followed a different distribution than those reported previously (Fendrych et al., 2013). This difference reflects methodological advances that improve the detection of short-lived events, which may have been underrepresented in earlier studies relying on manual tracking. Indeed, shorter exocytic dwell times have also been reported using more recent imaging approaches in plants (Lindeboom et al., 2026). However, recent studies in yeast have reported longer average dwell times, similar to those reported by Fendrych et al. (2013) (Kramer et al., 2026; Puig-Tintó et al., 2026). It is important to note that we included all detected EXO84b events and did not distinguish between bona fide and ambiguous exocytic events. Our observations raise the possibility that WAVE/SCAR–ARP2/3 contributes to localized exocytic events or post-fusion membrane organization. Although our data indicate a strong relationship between ARP2/3 and exocytosis, colocalization with TPLATE and dynamin was also significant. In non-plant organisms, the roles of the nucleation-promoting factor N-WASP (Benesch et al., 2005; Merrifield et al., 2004; Otsuki et al., 2003) and cortactin (Le Clainche et al., 2007) in activating the Arp2/3 complex at sites of clathrin-mediated endocytosis are well documented. In plant cells, current models indicate that actin is largely dispensable for the early stages of clathrin-mediated endocytosis (Narasimhan et al., 2020) and the functional interaction between the plant NPF WAVE/SCAR and endocytic processes has not yet been demonstrated. Therefore, the occasional proximity to endocytic markers such as TPLATE may reflect coupling of secretion and internalization events at the plant cortex (Yan et al., 2021; L. Zhang et al., 2019), rather than a direct involvement of ARP2/3 in endocytic pit formation. In plants, a link between ARP2/3 and endocytosis has been documented for the protein *At*EH1/Pan1, a subunit of the TPLATE complex involved in plant endocytosis (Wang et al., 2019). At EPCS, *At*EH1/Pan1 cooperates with VAP27, ARP2/3, and actin in autophagosome formation. It would be interesting to determine whether *At*EH1/Pan1 is involved in the formation of the ARP2/3 foci observed in our study and whether such a connection could provide further insight into the role of ARP2/3 in plant endocytosis. The specific targeting of ARP2/3 activity to distinct cellular locations supports a model in which the plant ARP2/3–WAVE/SCAR acts as a module that is activated only transiently and locally to fulfil diverse cellular functions. This spatially and temporally restricted organization of ARP2/3 activity may explain why mutants lacking ARP2/3 exhibit pronounced developmental phenotypes despite retaining largely intact bulk actin networks (Cifrová et al., 2020; García-González et al., 2026; Xu et al., 2024). In this view, ARP2/3 functions less as a global actin organizer and more as a dynamic cortical hub fine-tuning membrane trafficking at specific plasma membrane domains. Such a model provides a plausible explanation for the well-documented cell wall and growth defects of ARP2/3 and WAVE/SCAR mutants. In this work, we uncovered a highly dynamic population of the WAVE/SCAR–ARP2/3 module at the plasma membrane that is associated with exocytotic and endocytotic processes. In contrast, the accumulation of WAVE/SCAR at three-way junctions is highly stable, and given the inability to detect ARP2/3 at these sites, it is likely that WAVE/SCAR activates ARP2/3 in a very rapid and transient manner that precludes its detection by confocal microscopy. In both cases, the module is likely localized in close proximity to, or directly at, the plasma membrane. Indeed, several studies have reported plant ARP2/3 membrane association (Dyachok et al., 2008; Kotchoni et al., 2009). However, we cannot unequivocally confirm that the cortical foci are membrane-associated. The stable population of NAP1-GFP at the periphery of three-way cell junction may indicate a stable membrane localization.

However, we lack evidence for the mechanism by which WAVE/SCAR could be anchored at these sites. Our finding that ARP2/3 subunits co-immunoprecipitated with EXO84b, a subunit of the exocyst complex, provides one possible mechanism for the interaction of ARP2/3 with membranes, namely via the membrane-associated exocyst complex. However, as ARP2/3 dynamics and foci density remained unchanged in *exo84b* mutant plants, another exocyst subunit may serve as the anchoring component, such as EXO70A1. Indeed, EXO70A1 remains partially associated with the plasma membrane in the *exo84b* mutant (Synek et al., 2021). Interestingly, the co-immunoprecipitation of ARPC5 with EXO84b was consistently more efficient than that of ARPC2. It is possible that fluorescent tags influence the efficiency of interactions or that ARPC5, as a peripheral subunit, is more likely to engage in interactions with other proteins compared to the core subunit ARPC2.

In summary, we described a highly dynamic population of ARP2/3 complexes that colocalize with the cytoskeleton, colocalize with exocytic and endocytic markers in the cortical cytoplasm and co-immunoprecipitated with an exocyst protein subunit. Our work positions the plant WAVE/SCAR–ARP2/3 module within a dynamic cortical hub that links cytoskeletal regulation with spatially restricted membrane trafficking at the cell surface. We propose that the majority of the ARP2/3 in the cortical cytoplasm is maintained in an incomplete and thus inactive state, and that ARP2/3 is activated only locally.

## Acknowledgements

The work was supported by the grants of the Grant Agency of the Charles University no. 374522 (B.J., M.V. and K.S.) and TowArds Next GENeration Crops [reg. no. CZ.02.01.01/00/22_008/0004581] of the ERDF Programme Johannes Amos Comenius. Confocal and TIRF/VAE microscopy was performed in the Vinicna Microscopy Core Facility (RRID:SCR_026602) co-financed by the MEYS CR LM2023050 Czech-BioImaging. Computational resources were supplied by the e-INFRA CZ project (ID:90254) provided within the program Projects of Large Research, Development, and Innovations Infrastructures. Spinning disc microscopy and FCS was performed at the Imaging Facility of the Institute of Experimental Botany AS CR supported by the MEYS CR LM2023050 Czech-BioImaging, the Czech Academy of Sciences and IEB AS CR. Elyra 7 imaging was performed at the Light Microscopy Core Facility, IMG, Prague, Czech Republic, supported by MEYS CR LM2023050 Czech-BioImaging, MEYS – CZ.02.1.01/0.0/0.0/18_046/0016045 and MEYS – CZ.02.01.01/00/23_015/0008205.

The constructs 35S promoter, mScarlet-I and the *35S* terminator used for the GoldenBraid cloning were a gift from Matyáš Fendrych. We thank Prof. Fatima Cvrčková for critical reading of the manuscript. AI tools were used to facilitate translation and code writing.

## Competing interests

Authors declare no competing interests.

## Author contributions

B.J., V.Z., J.P., and K.S. contributed to the experimental design. B.J., M.V., K.L., J.K., J.G.-G., K.M., A.S., J.P., K.S. contributed to microscopy and image analysis. B.J., M.V., J.G.-G., S.V. and A.H. contributed to data analysis and statistics. B.J., M.V., E.K. and A.B.F. contributed to cloning and transformation of the plant lines. B.J. and K.S. contributed to Western blotting. B.J. and K.S. contributed to the writing of the manuscript. All authors red and approved the manuscript.

## Data availability

The data supporting the findings of this study are available from the corresponding authors upon request. Custom image analysis code developed for this study are available in public GitHub repositories.

## Supporting Information

**Figure S1.**

Detection of cortical ARP2/3 is strongly dependent on expression system. Analysis of the correlation between signal intensity and the dwell time. The FCS analysis shows a decreased diffusion coefficient for GFP-ARPC2 compared to free GFP. FRAP analysis of cytoplasmic diffuse signal and three-way junction NAP1-GFP accumulations. 3D reconstruction of pNAP1::GFP-NAP1 signal in 5DAS cotyledon epidermal cells.

**Figure S2.**

Comparison of XVE::GFP-ARPC2 dynamics and density in rescued (*arpc2*) and mutant (*arpc5*) lines. Supplemental plots for colocalization of 35S::mSc-ARPC5 and XVE::GFP-ARPC2 (D, F) and 35S::mSc-ARPC5 and pNAP1::NAP1-GFP.

**Figure S3.**

Supplemental information for image analysis of colocalization with the cytoskeleton.

**Figure S4.**

Analysis of cytoskeletal inhibitors CK666, BDM and PBP the dynamics and density of ARP2/3 cortical foci.

**Figure S5.**

Supplemental information for analysis of ARP2/3 foci relationship to membrane trafficking processes.

**Figure S6.**

Full blots and electrophoresis images for Fig. 5

**Figure S7.**

Phenotypes of plants of newly introduced lines.

**Table S1.**

Primers used for cloning.

**Table S2.**

Dwell time statistics for ARP2/3 and NAP1 foci.

**Table S3.**

Statistics for colocalization of ARPC5 with ARPC2 or NAP1.

**Table S4.**

Statistics for colocalization of *pUBQ::*GFP-ARPC5 with AFs or MTs.

**Table S5.**

Statistics for colocalization of ARP2/3 with markers for exocytosis and endocytosis.

**Video S1.**

ARP2/3 complex subunit ARPC2 localizes as dynamic foci in the cortical region of epidermal cell cytoplasm.

**Video S2.**

ARP2/3 complex subunit ARPC5 localizes as dynamic foci in the cortical region of epidermal cell cytoplasm.

**Video S3.**

WAVE/SCAR complex subunit NAP1 localizes as dynamic foci in the cortical region of epidermal cell cytoplasm.

**Video S4.**

3D reconstruction of pNAP1::GFP-NAP1 signal in 5DAS cotyledon epidermal cells.

**Video S5.**

3D reconstruction of pNAP1::GFP-NAP1 signal in 5DAS cotyledon epidermal cells in mutant background.

**Video S6.**

ARP2/3 subunit colocalizes with actin filaments.

**Video S7.**

ARP2/3 subunit colocalizes with microtubules.

## Notes

### Competing Interest Statement

The authors have declared no competing interest.

### Summary of Updates

This version has been revised to update the title, author list, and address several new findings in the field that were not mentioned in the previous version. The main figures are now also embedded in the text rather than at the end.

