## Supporting info for "*In vivo* imaging uncovers an abundant but rarely active pool of plant ARP2/3 and its link to exocytosis and endocytosis"

Article acceptance date: Click here to enter a date.

The following Supporting Information is available for this article:

**Fig. S1** Detection of cortical ARP2/3 is strongly dependent on expression system. Analysis of the correlation between signal intensity and the dwell time. The FCS analysis shows a decreased diffusion coefficient for GFP-ARPC2 compared to free GFP. FRAP analysis of cytoplasmic diffuse signal and three-way junction NAP1-GFP accumulations. 3D reconstruction of pNAP1::GFP-NAP1 signal in 5DAS cotyledon epidermal cells.

**Fig. S1:**

Detection of cortical ARP2/3 is strongly dependent on expression system. (a) 35S::GFP-ARPC2; 35S::GFP-ARPC5 and 35S::mSc-ARPC5 expressed in hypocotyl epidermal cells and *nap1*/pNAP1::NAP1-GFP expressed in cotyledon pavement cells (VAEM). Note increased diffuse signal and limited resolution of ARP2/3 foci expressed under 35S promoter. (b) Analysis of the correlation between signal intensity and the dwell time demonstrates that the signal mean intensity of foci does not correlate with their dwell time. Mean signal intensity of individual tracks (normalized by median value for each sample separately) is plotted against dwell times for *arpc5*/pUBQ::GFP-ARPC5 and *nap1*/pNAP1::NAP1-GFP. The top histogram shows dwell time distribution (count of events), the histogram on the right shows distribution of signal intensities normalized to the median value. (c) The FCS analysis shows a decreased diffusion coefficient (DC) for GFP-ARPC2 compared to free GFP, indicating reduced molecular mobility and suggesting that ARPC2 diffuses as part of a larger molecular complex. Diffusion coefficients are shown across the full range of measured FCS-fitted concentrations and induction conditions. Lines represent predictions from the concentration-adjusted linear mixed-effects model, with shaded areas indicating 95% confidence intervals. (d) FRAP analysis of cytoplasmic diffuse signal and three-way junction NAP1-GFP accumulations. While diffuse cytoplasmic signal recovered within 30 minutes after photobleaching, enriched NAP1-GFP signal in three-way junctions did not. Signal intensity was normalized to the signal intensity before bleaching. (e, f) 3D reconstruction of pNAP1::GFP-NAP1 signal in 5DAS cotyledon epidermal cells. (e) NAP1 localizes to three-way cell junctions (yellow arrow), accumulations at anticlinal cell walls, probably representing pit fields (Chi & Ambrose, 2025) (green arrow), and associates with peroxisomes (magenta arrow) in *nap1* rescued line. (f) The same marker expressed in *brick1* mutant background fails to localize to three-way junctions (yellow arrow) and anticlinal cell walls and remains associated only with peroxisomes (magenta arrow). For more detail see Supplemental Videos S4, 5. Brightness and contrast have been modified for images (a, e, and f) for representative reasons. Scale bar: 5 µm. Side of one square in the grid box in images (e, f) is
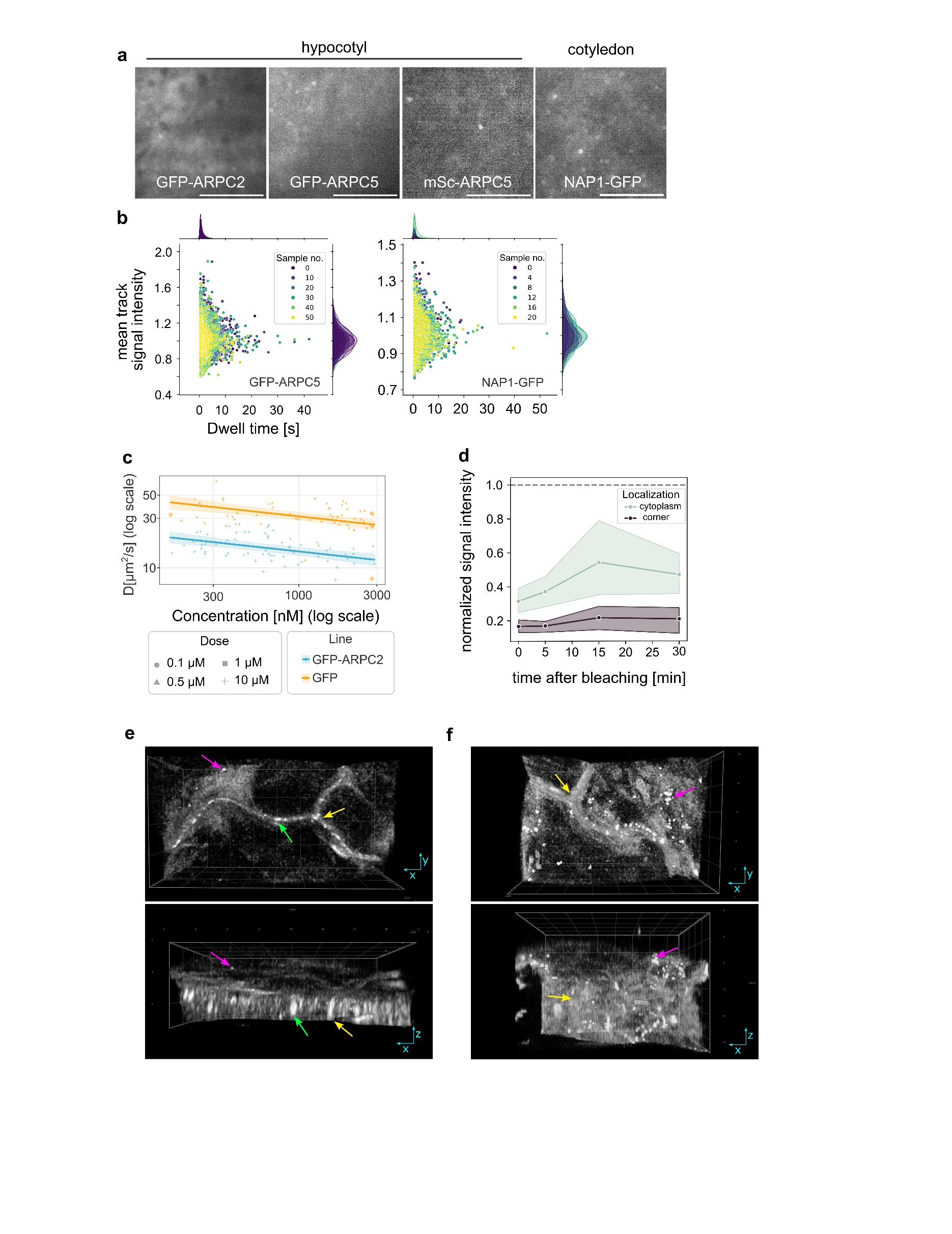
4 µm.

**Fig. S2** Comparison of XVE::GFP-ARPC2 dynamics and density in rescued (*arpc2*) and mutant (*arpc5*) lines. Supplemental plots for colocalization of 35S::mSc-ARPC5 and XVE::GFP-ARPC2 (D, F) and 35S::mSc-ARPC5 and pNAP1::NAP1-GFP.

**Fig. S2:**

(a, b) Comparison of XVE::GFP-ARPC2 dynamics and density in rescued (*arpc2*) and mutant (*arpc5*) lines. (a) *arpc5*/XVE::GFP-ARPC2 (VAEM, hypocotyl epidermal cell). Asterisks label previously reported ARP2/3 peroxisomal localization; this area was omitted from all the analyses. (b) Proportion of foci with certain dwell time (binned) of XVE::GFP-ARPC2 foci is comparable in *arpc2* and *arpc5* mutants, although follows the same trend as seen in GFP-ARPC5 foci expressed in *arpc2*. (c) Comparison of density of XVE::GFP-ARPC2 in *arpc2* (rescue) or *arpc5* mutant background. Number of ARPC2 foci is significantly lower in *arpc5* mutant (p < 0.0001). (d, e) Analysis of colocalization of 35S::mSc-ARPC5 and XVE::GFP-ARPC2 (d, f) and 35S::mSc-ARPC5 and pNAP1::NAP1-GFP (d, f). (d) VAEM in hypocotyl epidermal cells. (e) Percentage of mSc-ARPC5 foci colocalizing with GFP-ARPC2 (magenta; p < 0.0001) and percentage of GFP-ARPC2 foci colocalizing with mSc-ARPC5 (green) compared to negative controls (90 ° rotated image; p < 0.0001). (f) Percentage of mSc-ARPC5 foci colocalizing with NAP1-GFP (magenta) and percentage of NAP1-GFP foci colocalizing with mSc-ARPC5 (green; p < 0.0001) compared to negative controls (90 ° rotated image; p < 0.05). Note that the proportion of colocalizing foci is higher for mSc-ARPC5 because it contains fewer total foci than GFP-ARPC2 or NAP1-GFP, while the number of colocalizing foci remains unchanged. (g) Plots showing foci/background signal intensity ratio for XVE:GFP-ARPC2 and pUBQ::GFP-ARPC5 in *arpc2* or *arpc5* mutant backgrounds (p < 0.01 and p <0.0001 respectively). Note improved foci/background signal intensity ratio in rescued mutant lines.

Statistical analysis: bootstrapping – error bars around timepoint values represent the 95% confidence intervals - if not overlapping, the result is considered statistically significant (b); Welch's t-test independent samples (c); Wilcoxon test (e, f); t-test independent samples (g). Brightness and contrast have been modified for images (a) and (d) for illustrative reasons. Scalebar: 5 µm.

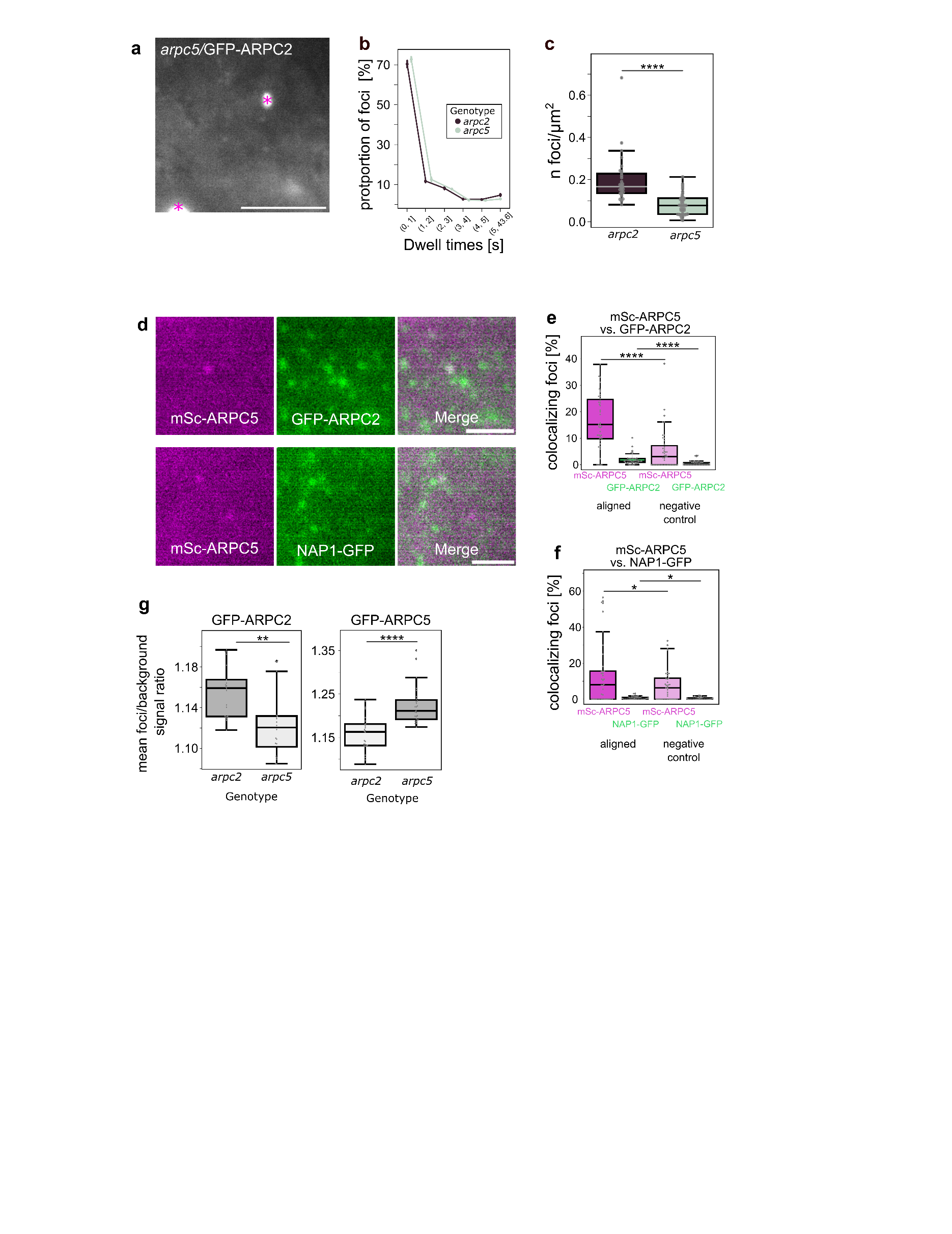

**Fig. S3** Supplemental information for image analysis of colocalization with the cytoskeleton.

**Fig. S3:**

Image analysis of colocalization with the cytoskeleton. (a) Example of a mask for actin filaments used for colocalization; (b) example of a mask for microtubules used for colocalization. (c) Occasional reorganization of GFP-ARPC5 foci into a pattern resembling microtubules (arrows) upon LatB treatment of pUBQ::GFP-ARPC5 expressing hypocotyl pavement cells. (d, e) Representative VAEM images of cortical cytoplasm of hypocotyl pavement cells of *arpc5*/pUBQ::GFP-ARPC5 line untreated and treated with latrunculin B (d) and oryzalin (e). Asterisks label previously reported ARP2/3 peroxisomal localization. Brightness and contrast in images have been modified for representative reasons. Scalebar: 5 µm

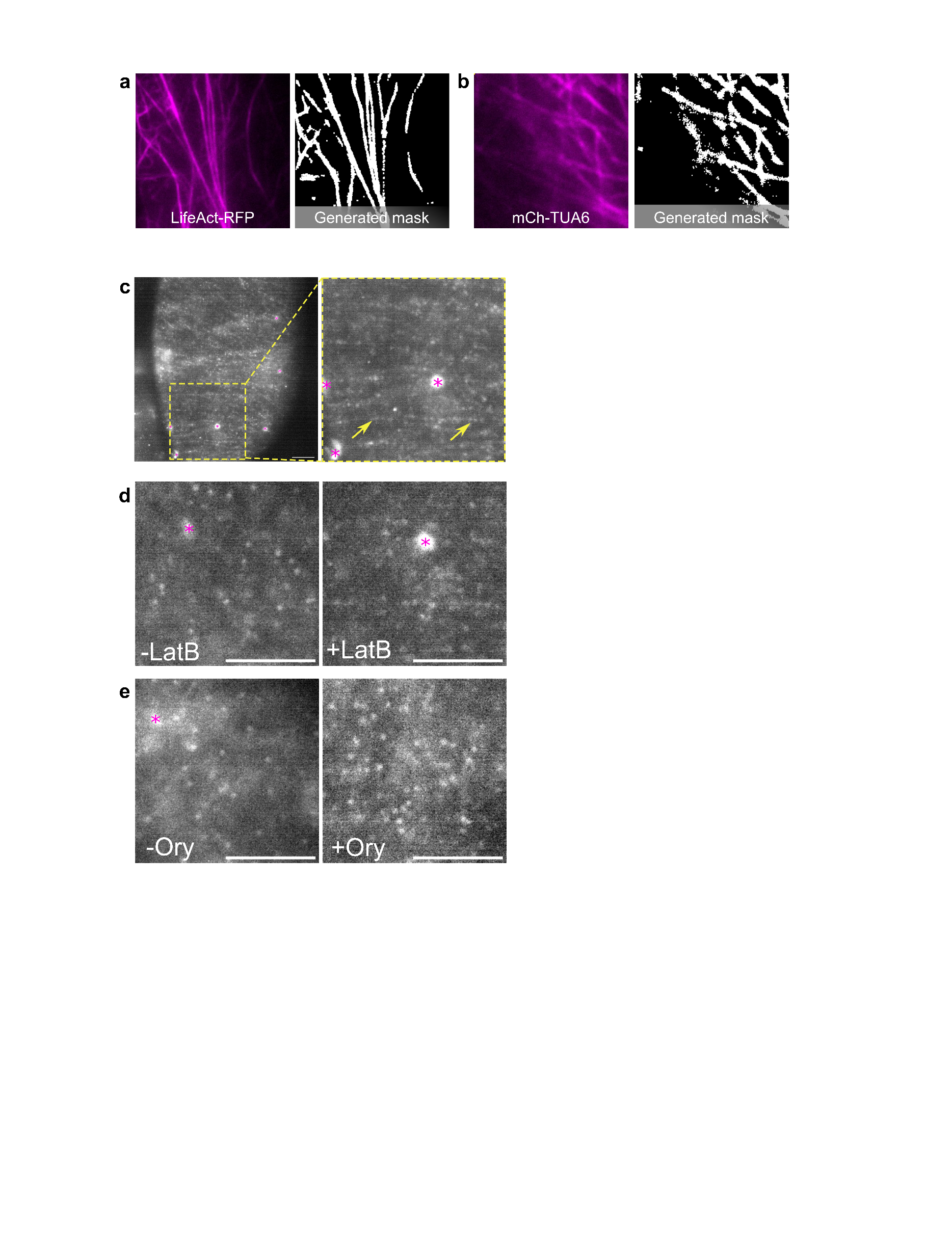

**Fig. S4** Analysis of cytoskeletal inhibitors CK666, BDM and PBP the dynamics and density of ARP2/3 cortical foci.

**Fig. S4:**

Analysis of cytoskeletal inhibitors CK666 (a-c), BDM (d-f) and PBP (g-i) the dynamics and density of ARP2/3 cortical foci. (a, d, g) *arpc5*/pUBQ::GFP-ARPC5 line treated with ARP2/3 inhibitor CK666 (5 min, 10 µM) (a), myosin inhibitor BDM (30 min, 50 mM) (d), and myosin inhibitor PBP (5 min, 10 µM) (g) with respective mock-treated control. Hypocotyl epidermal cells imaged by VAEM; asterisks label previously reported ARP2/3 peroxisomal localization; this area was omitted from all the analyses. (b, e, h) Analysis of dynamics of GFP-ARPC5 foci shown as a proportion of foci with certain dwell time (binned) in CK666 (b), BDM (e) and PBP (h) treated cells compared to respective control demonstrated unchanged dwell times. (c, f, i) Analysis of density of GFP-ARPC5 foci in CK666 (c), BDM (f) and PBP (Ii treated cells compared to respective control demonstrated unchanged density in treated cells. Statistical analysis: bootstrapping – the result is considered statistically significant if error bars around timepoint values representing the 95% confidence intervals do not overlap (b, e, h); Welch's t-test for independent samples (c, f, i); p values > 0.05 (not significant, ns). Brightness and contrast in images (a-f) have been modified for representative reasons. Scalebar: 5 µm.

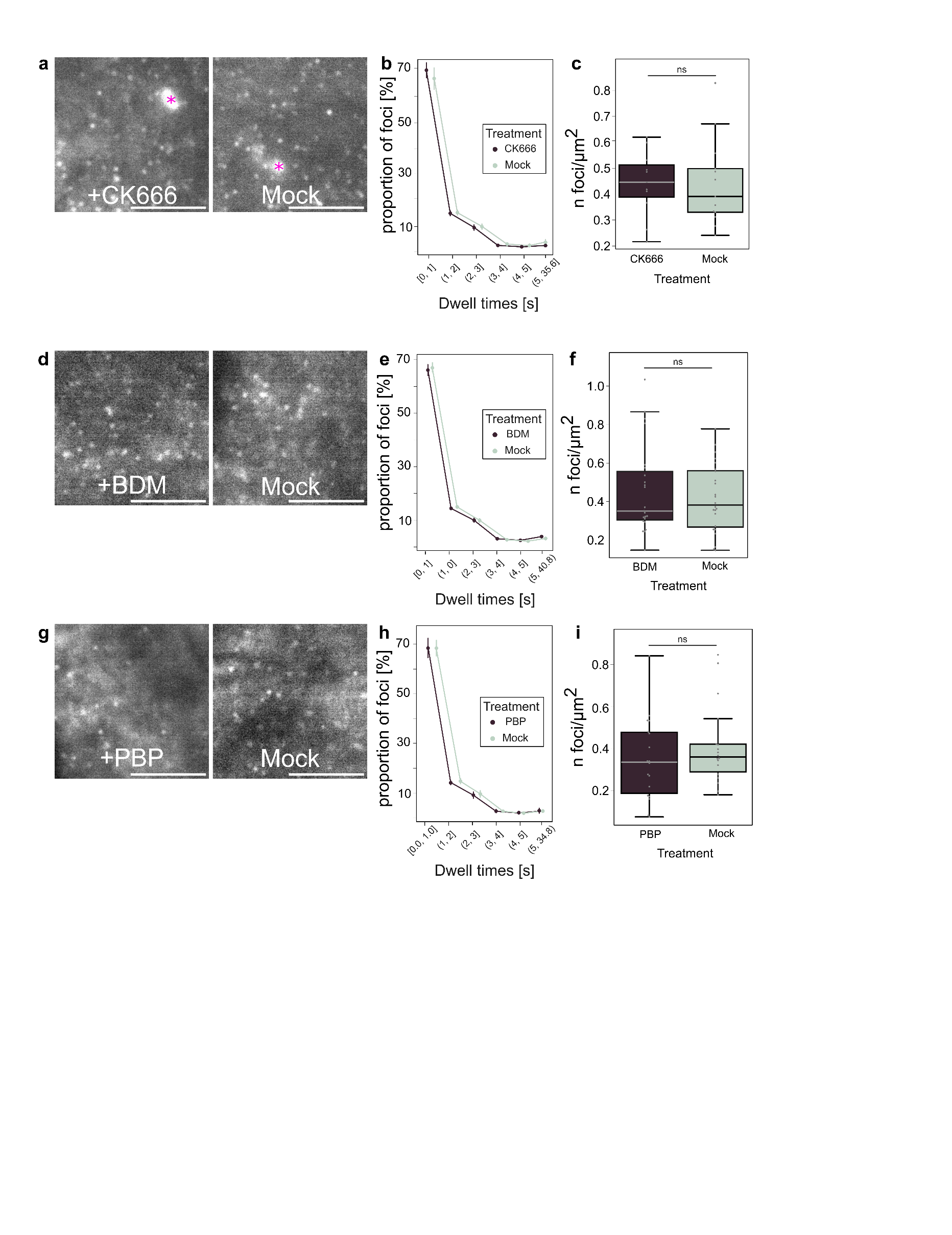

**Fig. S5** Supplemental information for analysis of ARP2/3 foci relationship to membrane trafficking processes.

**Fig. S5:**

ARP2/3 foci relationship to membrane trafficking processes. (a, b) Dynamics and density of ARP2/3 foci is unchanged in *exo84b*. XVE::GFP-ARPC2 foci dwell times are comparable in *exo84b* and wt plants, as the proportion of foci with certain dwell time (binned) is statistically not significant (a). Density of XVE::GFP-ARPC2 cortical foci does not differ significantly in *exo84b* and wt plants (p > 0.05) (b). (c, d) Analysis of ARP2/3 markers XVE::GFP-ARPC2 and 35S::GFP-ARPC5 and membrane trafficking markers 35S::EXO84b-RFP and 35S::TPLATE-RFP colocalization calculated reciprocally for both markers combinations (see Fig. 4b, c for calculation of ARP2/3 colocalization with membrane trafficking markers only). (c) Percentage of GFP-ARPC2 foci colocalizing with EXO84b-RFP compared to negative controls (90 ° rotated image; p < 0.0001, green) and percentage of EXO84b-RFP foci colocalizing with GFP-ARPC2 compared to negative controls (90 ° rotated image; p < 0.0001, magenta). (d) Percentage of GFP-ARPC5 foci colocalizing with TPLATE-RFP compared to negative controls (90 ° rotated image; p < 0.01, green) and percentage of TPLATE-RFP foci colocalizing with GFP-ARPC5 compared to negative controls (90 ° rotated image; p < 0.01, magenta). (e, f) Analysis of colocalization of XVE::GFP-ARPC2 and pDRP1C::DRP1C-mOrange. (e) VAEM in hypocotyl epidermal cells. (f) Percentage of GFP-ARPC2 foci colocalizing with DRP1C-mOra compared to negative controls (90 ° rotated image; p < 0.001, green) and percentage of DRP1C-mOra foci colocalizing with GFP-ARPC2 compared to negative controls (90 ° rotated image; p < 0.01, magenta).

Note that the proportion of colocalizing foci is higher for the marker with fewer total foci, although the absolute number of colocalizing foci remains the same.

Statistical analysis: bootstrapping - lines around timepoint values represent the 95% confidence intervals - if not overlapping, the result is considered statistically significant (a); Welch's t-test for independent samples (b); Wilcoxon test (c, d, f).

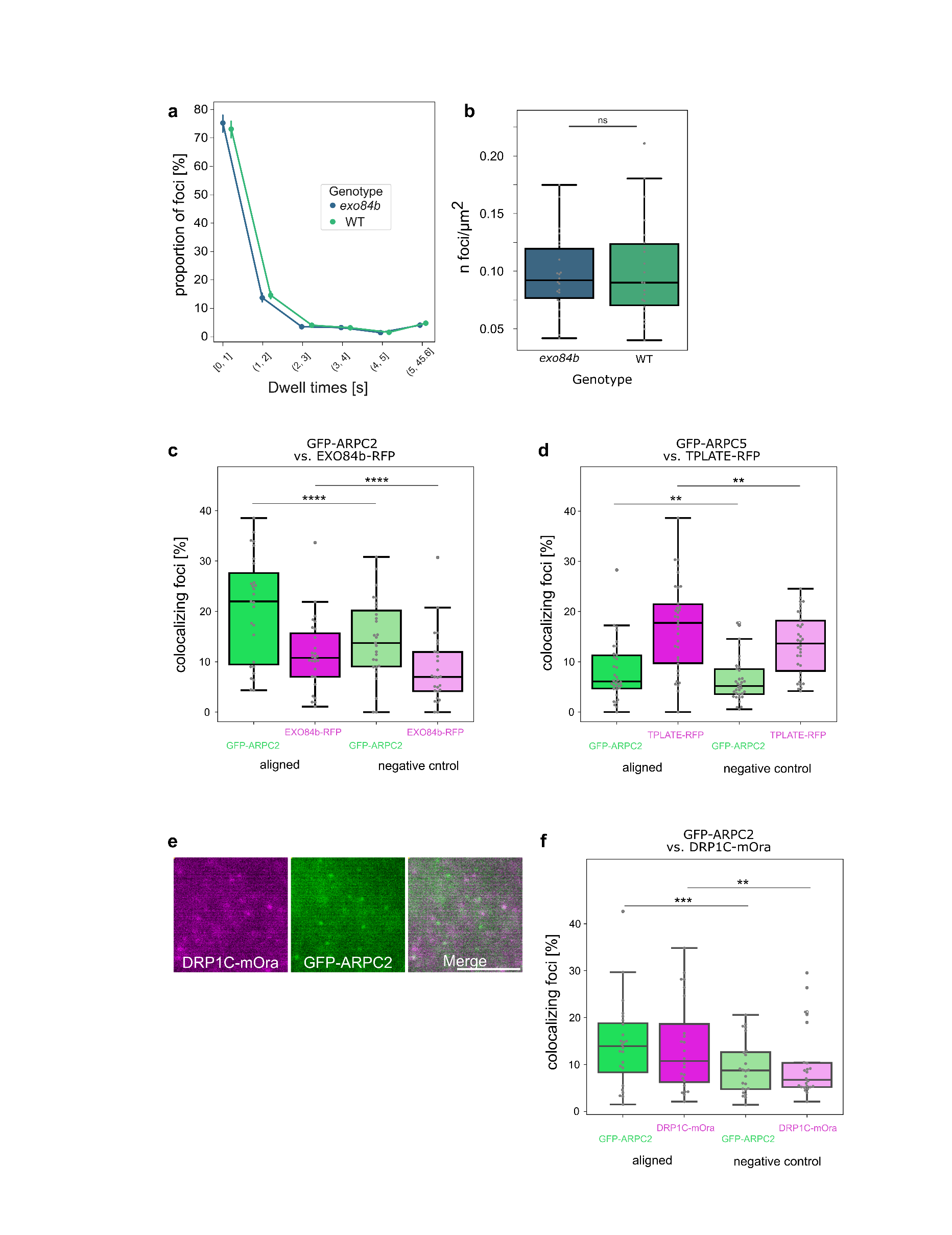

**Fig. S6** Full blots and electrophoresis images for Fig. 5

**Fig. S6:**

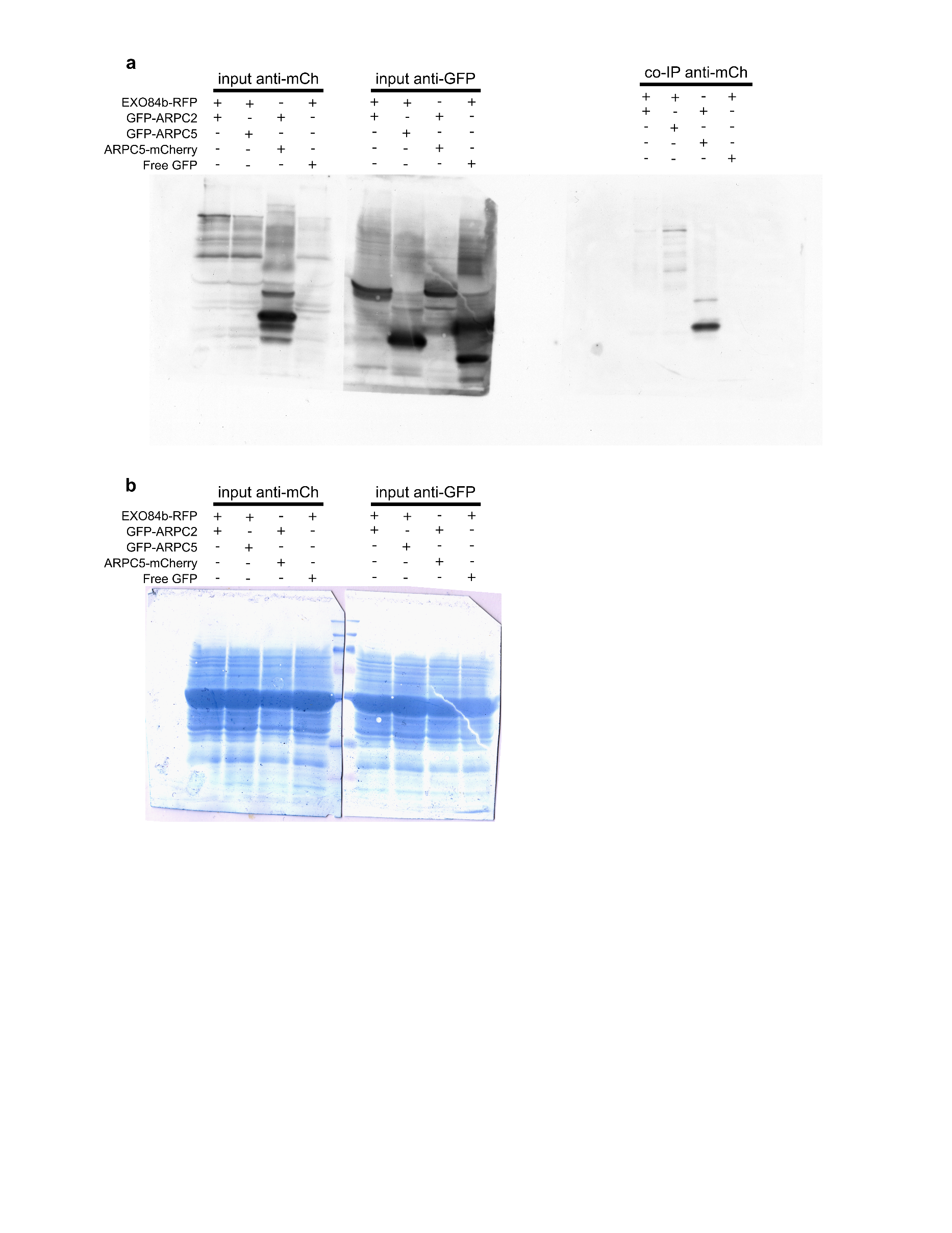
Full blots and electrophoresis images for Fig. 5- biochemical analysis of EXO84b interaction with ARP2/3. (a) protein extracts of inputs for co-IP and co-IP probed with anti-mCherry and anti-GFP antibodies. (b) amido black-stained (0.1% (w/v) Amido Black 10B I 10%(v/v) and acetic acid and 45% ethanol, 5 minute incubation) membrane with inputs to demonstrate equal gel loading.

**Fig. S7** Phenotypes of plants of newly introduced lines.

**Fig. S7:**

ARP2/3 markers for ARPC5 subunit used in this study rescue ARP2/3 mutant phenotype of distorted trichomes (a, b). Expression of different ARP2/3 complex subunit does not rescue the mutant phenotype (c, d). Likewise, expression of an ARP2/3 complex subunit does not rescue the phenotype of a mutant lacking a WAVE/SCAR complex subunit (e).

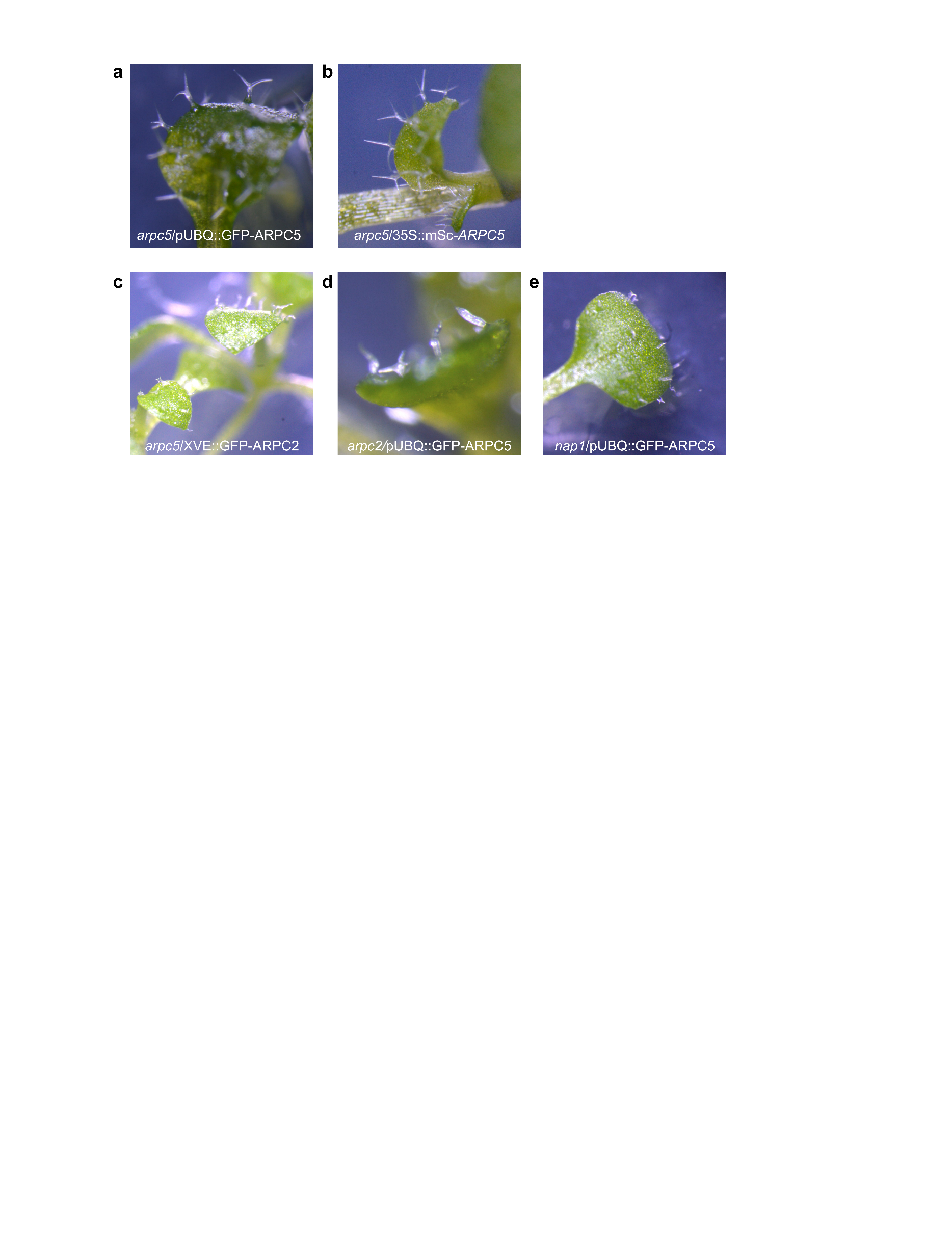

**Table S1** Primers used for cloning.

| For amplification of ARPC5 CDS from vector DNA for the 35S::mSc-ARPC5 | forward - GCGCCGTCTCGCTCGAATGGCAGAATTCGTTGAAGCT |
| --- | --- |
|  | reverse - CGCCGTCTCGCTCAAAGCTCAAACGGTGTTGATGGTATCAGTAAGACATCGGAGAATGCAACCAAGACCAGCT |
| For amplification of AtARPC5 genomic sequence, including the stop codon for the pUBQ::GFP-ARPC5 | forward - GGGGACAGCTTTCTTGTACAAAGTGGCTATGGCAGAATTCGTTGAAGCTG |
|  | reverse -GGGGACAACTTTGTATAATAAAGTTGCTCAAACGGTGTTGATGGTATCAG |

**Table S2** Dwell time statistics for ARP2/3 and NAP1 foci (Fig. 1b).

|  | *arpc5/*pUBQ::GFP-ARPC5 | *arpc2/*XVE*:*:GFP-ARPC2 | *nap1/*pNAP1::GFP-NAP1 |
| --- | --- | --- | --- |
| mean [sec] | 1.091330 | 1.264625 | 1.153394 |
| std [sec] | 1.401469 | 2.006725 | 2.006725 |
| min [sec] | 0.400016 | 0.400016 | 0.400016 |
| median [sec] | 0.800029 | 0.400016 | 0.400016 |
| max [sec] | 42.001678 | 43.601872 | 52.801869 |
| n events | 40352 | 38907 | 28948 |
| n ROIs | 51 | 28 | 19 |

**Table S3** Statistics for colocalization of ARPC5 with ARPC2 or NAP1 (Fig. 2d, h; Figure S2d-f).

|  | ARPC2 x ARPC5  aligned | ARPC2 x ARPC5  control | ARPC5 x ARPC2  aligned | ARPC5x ARPC2  control | NAP1 x ARPC5 aligned | NAP1 x ARPC5  control | ARPC5 x NAP1 aligned | ARPC5 x NAP1 control |
| --- | --- | --- | --- | --- | --- | --- | --- | --- |
| min [%] | 0 | 0 | 0 | 0 | 0 | 0 | 0 | 0 |
| mean [%] | 1,86 | 0,59 | 16,54 | 5,36 | 0,74 | 0,49 | 14,03 | 7,85 |
| max [%] | 10,13 | 3,46 | 37,84 | 38,1 | 3,09 | 2,17 | 56,52 | 32,43 |
| n events analyzed | 18660 | 18657 | 1631 | 1631 | 22064 | 22073 | 1270 | 1270 |
| n ROIs analyzed | 43 | 43 | 43 | 43 | 43 | 38 | 38 | 38 |

**Table S4** Statistics for colocalization of *pUBQ::*GFP-ARPC5 with AFs or MTs (Fig. 3b, c) .

|  | Coloc with AFs | control coloc with AFs | Coloc with MTs | control coloc with MTs |
| --- | --- | --- | --- | --- |
| min [%] | 27,42 | 24,08 | 46,48 | 51,31 |
| mean [%] | 46,32 | 37,93 | 73,01 | 64,12 |
| max [%] | 78,92 | 60,87 | 83,56 | 73,16 |
| n events analyzed | 21705 | 21705 | 15898 | 15898 |
| n ROIs analyzed | 29 | 29 | 32 | 32 |

**Table S5** Statistics for colocalization of ARP2/3 with markers for exocytosis and endocytosis (Figure 4b, c; Fig. S5c, e, f).

|  | ARP x EXO aligned | ARP x EXO  control | EXO x ARP  aligned | EXO x ARP  control | ARP x TPLATE aligned | ARP x TPLATE  control | TPLATE x ARP aligned | TPLATE x ARP control | ARP x  DRP  aligned | ARP x  DRP  control | DRP x  ARP aligned | DRP x  ARP control |
| --- | --- | --- | --- | --- | --- | --- | --- | --- | --- | --- | --- | --- |
| min [%] | 4.35 | 1.09 | 0.00 | 0.00 | 0.00 | 0.54 | 0.00 | 4.17 | 1.49 | 1.42 | 2.11 | 2.11 |
| mean [%] | 20.62 | 11.41 | 14.43 | 8.51 | 8.03 | 6.35 | 16.85 | 13.50 | 14.22 | 9.36 | 13.79 | 9.98 |
| max [%] | 38.53 | 33.66 | 30.80 | 30.69 | 28.28 | 17.73 | 38.62 | 24.54 | 42.64 | 20.59 | 34.85 | 29.56 |
| n events analyzed | 7721 | 7793 | 14716 | 14716 | 23593 | 23593 | 9210 | 9246 | 4853 | 4877 | 5217 | 5217 |
| n ROIs analyzed | 27 | 27 | 27 | 27 | 33 | 33 | 33 | 33 | 24 | 24 | 24 | 24 |

**Video S1** ARP2/3 complex subunit ARPC2 localizes as dynamic foci in the cortical region of epidermal cell cytoplasm.

**Video S1:**

*arpc2/*XVE::GFP-ARPC2 in 5DAS hypocotyl epidermal cell. Visualized by VAEM, exposure time 400 ms. Brightness and contrast were adjusted for illustrative purposes. Scale bar: 5 µm, time: mm:ss.

**Video S2** ARP2/3 complex subunit ARPC5 localizes as dynamic foci in the cortical region of epidermal cell cytoplasm.

**Video S2:**

*arpc5/*pUBQ::GFP-ARPC5 in 5DAS hypocotyl epidermal cell. Visualized by VAEM, exposure time 400 ms. Brightness and contrast were adjusted for illustrative purposes. Scale bar: 5 µm, time: mm:ss.

**Video S3** WAVE/SCAR complex subunit NAP1 localizes as dynamic foci in the cortical region of epidermal cell cytoplasm.

**Video S3:**

*nap1/*pNAP1::NAP1-GFP in 5DAS hypocotyl epidermal cell. Visualized by VAEM, exposure time 400 ms. Brightness and contrast were adjusted for illustrative purposes. Scale bar: 5 µm. time: mm:ss.

**Video S4** 3D reconstruction of pNAP1::GFP-NAP1 signal in 5DAS cotyledon epidermal cells.

**Video S4:**

NAP1 localizes to three-way cell junctions (yellow arrow), accumulations at anticlinal cell walls, probably representing pit fields (Chi & Ambrose, 2025) (green arrow), and associates with peroxisomes (Martinek et al., 2023) (magenta arrow) in *nap1* rescued line. Visualized by laser scanning confocal microscope Leica SP8 equipped with HC PL APO CS2 63×/1.2 W objective. Brightness and contrast were adjusted for illustrative purposes. Side of one square in the grid box is 4 µm.

**Video S5** 3D reconstruction of pNAP1::GFP-NAP1 signal in 5DAS cotyledon epidermal cells in mutant background.

**Video S5:**

NAP1 marker expressed in *brick1* mutant background fails to localize to three-way junctions (yellow arrow) and anticlinal cell walls and remains associated only with peroxisomes (Martinek et al., 2023) (magenta arrow). Visualized by laser scanning confocal microscope Leica SP8 equipped with HC PL APO CS2 63×/1.2 W objective. Brightness and contrast were adjusted for illustrative purposes. Side of one square in the grid box is 4 µm.

**Video S6** ARP2/3 subunit colocalizes with actin filaments.

**Video S6:**

pUBQ::GFP-ARPC5 (green) and LifeAct-RFP (magenta) in 5DAS hypocotyl epidermal cell. Visualized by VAEM, exposure time 400 ms. Brightness and contrast were adjusted for illustrative purposes. Scale bar: 5 µm, time: mm:ss.

**Video S7** ARP2/3 subunit colocalizes with microtubules.

**Video S7:**

pUBQ::GFP-ARPC5 (green) and mCh-TUA5 (magenta) in 5DAS hypocotyl epidermal cell. Visualized by VAEM, exposure time 400 ms. Brightness and contrast were adjusted for illustrative purposes. Scale bar: 5 µm, time: mm:ss.
