## Supplemental Videos legends for "*In vivo* imaging uncovers an abundant but rarely active pool of plant ARP2/3 and its link to exocytosis and endocytosis"

Chi, Z., & Ambrose, C. (2025). The arabidopsis WAVE/SCAR protein BRICK1 associates with cell edges and plasmodesmata. *PLOS ONE*, *20*(9), e0325015. https://doi.org/10.1371/JOURNAL.PONE.0325015

Martinek, J., Cifrová, P., Vosolsobě, S., García-González, J., Malínská, K., Mauerová, Z., Jelínková, B., Krtková, J., Sikorová, L., Leaves, I., Sparkes, I., & Schwarzerová, K. (2023). ARP2/3 complex associates with peroxisomes to participate in pexophagy in plants. *Nature Plants*, *9*(11), 1874–1889. https://doi.org/10.1038/s41477-023-01542-6
